# Aiolos modulates the T_FH_ and CD4-CTL differentiation programs via reciprocal regulation of the Zfp831/TCF-1/Bcl-6 axis and CD25

**DOI:** 10.1101/2022.05.18.492485

**Authors:** Kaitlin A. Read, Devin M. Jones, Srijana Pokhrel, Emily D.S. Hales, Aditi Varkey, Jasmine A. Tuazon, Caprice D. Eisele, Omar Abdouni, Abbey Saadey, Robert T. Warren, Michael D. Powell, Jeremy M. Boss, Emily A. Hemann, Jacob S. Yount, Hazem E. Ghoneim, Chan-Wang J. Lio, Aharon G. Freud, Patrick L. Collins, Kenneth J. Oestreich

## Abstract

Effective immunity to influenza virus and other respiratory viruses requires the generation of CD4^+^ T cell subsets that coordinate multiple aspects of the immune response. These subsets include T follicular helper (T_FH_) and T helper 1 (T_H_1) cells, which promote humoral and cell-mediated responses, respectively. A third population, CD4^+^ cytotoxic T lymphocytes (CD4-CTLs) facilitates clearance of infection via mechanisms normally associated with CD8^+^ T cells. Here, we identify the transcription factor Aiolos as a regulator of T_FH_ and CD4-CTL responses. We demonstrate that Aiolos deficiency compromises T_FH_ differentiation and antibody production during influenza virus infection. Conversely, we find that CD4^+^ T cells acquire a cytotoxic-like program in the absence of Aiolos, including increased expression of the CTL-associated transcription factors Eomes and Blimp-1. We further show that while Aiolos positively regulates the T_FH_ transcriptional regulators Zfp831, TCF-1 and Bcl-6, it also directly represses expression of IL-2Rα and IL-2/STAT5-driven expression of the cytotoxic gene program. Thus, our findings identify Aiolos as a pivotal regulator of T_FH_ and CD4-CTL differentiation and highlight its potential as a target for manipulating CD4^+^ T cell humoral and cytotoxic responses.

## Introduction

Immune responses mounted against intracellular pathogens, such as influenza virus, require coordination between numerous immune cell types. These include the effector CD4^+^ T helper 1 (T_H_1), T follicular helper (T_FH_), and cytotoxic subsets, each of which perform distinct activities to clear infection. During infection, T_H_1 cells produce effector cytokines such as interferon gamma (IFN-γ) to both recruit and activate additional immune cell populations to sites of infection^1^. T_FH_ populations are key participants in the generation of humoral immune responses through their cognate B cell interactions, which support somatic hypermutation, class-switch recombination, affinity maturation, and the formation of long-lived plasma cell populations^2^. While CD4^+^ T cells have classically been appreciated as orchestrators of immune responses, it is now widely accepted that CD4^+^ T cells are also capable of differentiating into cytotoxic ‘CD4-CTLs’. CD4-CTLs are characterized by their ability to secrete both cytotoxic molecules and pro-inflammatory cytokines, ultimately resulting in the direct targeting and killing of virus-infected cells in a class two major histocompatibility complex (MHC-II) restricted fashion^3,4^. Together, the T_H_1, T_FH_, and CD4-CTL subsets provide the CD4^+^ T cell responses important for the clearance of infection. Their dysregulation has also been associated with autoimmune disorders in which inappropriate production of pro-inflammatory molecules and/or the production of autoantibodies drive pathogenesis^4–6^. Thus, due to their central roles in healthy and dysregulated immune responses, understanding the mechanisms that govern the differentiation and function of T_H_1, T_FH_, and CD4-CTL populations is of critical importance.

CD4^+^ T cell differentiation is initiated by antigen-presenting cells, which provide antigen-induced signals to the T cell receptor, as well as co-stimulation. Distinct polarization programs are then specified by cytokine signals, which induce the activity of a cascade of downstream transcriptional networks that ultimately imprint the unique transcriptional programs underlying each cell subset^7^. For example, IL-12/STAT4 signaling promotes the expression and function of the T-box factor T-bet, the T_H_1 lineage-defining transcription factor^8,9^. T-bet directly induces expression of IFN-γ, which positively regulates T-bet transcription in a feed-forward loop^10^. T_H_1 differentiation also requires the transcription factor Blimp-1, which represses expression of alternative polarization programs including those underlying the T_FH_ subset^11,12^. It is known that the T_H_1 and CD4-CTL gene programs share some regulatory features, including dependence on Blimp-1^4,13^. Furthermore, similar to T-bet in T_H_1 cells, CD4-CTL genes are positively regulated by the related T-box factor Eomesodermin (Eomes). However, to date, much of the transcriptional network underlying CD4-CTL populations remains undefined^4^.

Unlike T_H_1 and CD4-CTL populations, T_FH_ cell differentiation requires signaling from STAT3 activating-cytokines, such as IL-6, which induce the expression and function of the transcriptional repressor B cell lymphoma 6 (Bcl-6)^14^. Bcl-6 supports T_FH_ differentiation most notably by inhibiting the expression of the T_FH_ antagonist Blimp-1, as well as genes associated with alternative cells fates, including those important for T_H_1, CD4-CTL, T_H_2, and T_H_17 differentiation^11,15–17^. While Bcl-6 is considered the lineage-defining transcription factor for T_FH_ development, T_FH_ differentiation also relies upon TCF-1 (encoded by the gene *Tcf7*), Lef-1, Zpf831, Tox, Tox2, Maf, Batf, Ascl2, and Irf4^5,18^. Of these, Tox2, Batf, TCF-1, and Zfp831 have been implicated in the direct induction of *Bcl6* expression^19–24^. Like Bcl-6, TCF-1 also represses the expression of Blimp-1, and has been implicated in the negative regulation of both T_H_1 and CD4-CTL differentiation^15,19,23,25^. While much remains to be explored regarding the transcription factor Zfp831, it was recently identified as an upstream, positive regulator of both *Tcf7* and *Bcl6*^21^. Together, these early factors both set the stage for T_FH_ differentiation, and function to oppose alternative gene programs.

IL-2 signaling, which is propagated through tyrosine phosphorylation-dependent activation of STAT5, is also a key determinant in the regulation of T_H_1, T_FH_, and CD4-CTL populations. IL-2/STAT5 signaling supports T_H_1 and CD4-CTL generation, while opposing T_FH_ differentiation^3,4,12,26–29^. In T_H_1 and CD4-CTL populations, STAT5 directly induces expression of Blimp-1, and competes with the pro-T_FH_ transcription factor STAT3 to directly repress expression of *Bcl6* and additional T_FH_ gene targets^12^. During T_FH_ differentiation, IL-6 and IL-21-driven STAT3 represses IL-2 receptor expression and induces the expression of T_FH_ genes including Bcl-6^30^. Collectively, the available literature suggests that opposing sets of environmental and transcriptional regulators function to maintain the appropriate balance of T_H_1, T_FH_, and CD4-CTL effector responses during infection. Given the importance of this balance in maintaining health versus disease, the identity of factors that comprise these networks, as well as the unifying regulators that govern these relationships warrant further investigation.

Here, we have identified the Ikaros zinc finger transcription factor Aiolos as a central regulator of CD4^+^ T cell differentiation and effector responses to infection. We find that Aiolos deficiency cripples both T_FH_ cell differentiation and antibody production during influenza virus infection. We further find that loss of Aiolos allows for acquisition of a cytotoxic program that includes increased expression of key CTL transcriptional regulators and effector molecules. Mechanistically, we demonstrate that Aiolos positively regulates the expression of key T_FH_ transcription factors, while repressing IL-2Rα (CD25), through association with their respective gene regulatory elements. Consequently, Aiolos suppresses IL-2/STAT5-driven expression of the cytotoxic gene program. Together, these findings identify Aiolos as a reciprocal regulator of T_FH_ and CD4-CTL differentiation and support its potential as a target for the therapeutic manipulation of CD4^+^ T cell-dependent humoral and cytotoxic responses.

## Results

### Aiolos-deficient mice display defects in T_FH_ differentiation

In a prior study, we demonstrated that Aiolos cooperates with STAT3 to directly induce Bcl-6 expression in CD4^+^ T cells^31^. Given that Bcl-6 is the lineage-defining transcription factor for T_FH_ cells, we hypothesized that Aiolos is required for T_FH_ cell differentiation. To explore this possibility *in vivo*, we employed a well-established murine model of influenza virus infection. We began by infecting wildtype (WT) C57BL/6J mice with influenza A/PR/8/34 (H1N1), commonly termed PR8. We first evaluated Aiolos expression in naive CD4^+^ T cells, PD-1^+^Cxcr5^+^ T_FH_ cells, and non-TFH populations in lung-draining lymph nodes (DLN) (**Fig. 1A** and **Extended Data Fig. 1A**). Consistent with a role for Aiolos in T_FH_ populations generated *in vivo*, Aiolos protein expression was significantly elevated in T_FH_ populations relative to naive and non-TFH cells. Further, we found that Aiolos was highest in germinal center (GC) T_FH_ cells, characterized by high expression of PD-1 and Cxcr5 (**Fig. 1A**). To determine whether Aiolos is similarly increased in human T_FH_ cell populations, we analyzed previously published RNA-seq data for human naive, non-TFH, and T_FH_ cell populations from human tonsils (GSE58597)^32^. Indeed, human Aiolos *(IKZF3*) expression was highest in T_FH_ populations, and its expression correlated with that of key T_FH_ transcriptional regulators *(BCL6, TOX, TOX2*) and cell surface receptors (*CD40LG, PDCD1*, and *IL6R*) (**Extended Data Fig. 1B,C**). Expression of genes associated with antagonism of T_FH_ differentiation, including *PRDM1* (which encodes BLIMP-1) and *IL2RA*, inversely correlated with Aiolos expression. We next examined Aiolos protein expression in T_FH_ and non-TFH effector cells isolated from human tonsils. Again, Aiolos expression was elevated in T_FH_ populations relative to non-TFH effector cells (**Extended Data Fig. 1D,E**). Together, these findings supported a conserved role for Aiolos as a potential positive regulator of T_FH_ differentiation in both mice and humans.

**Figure 1.**
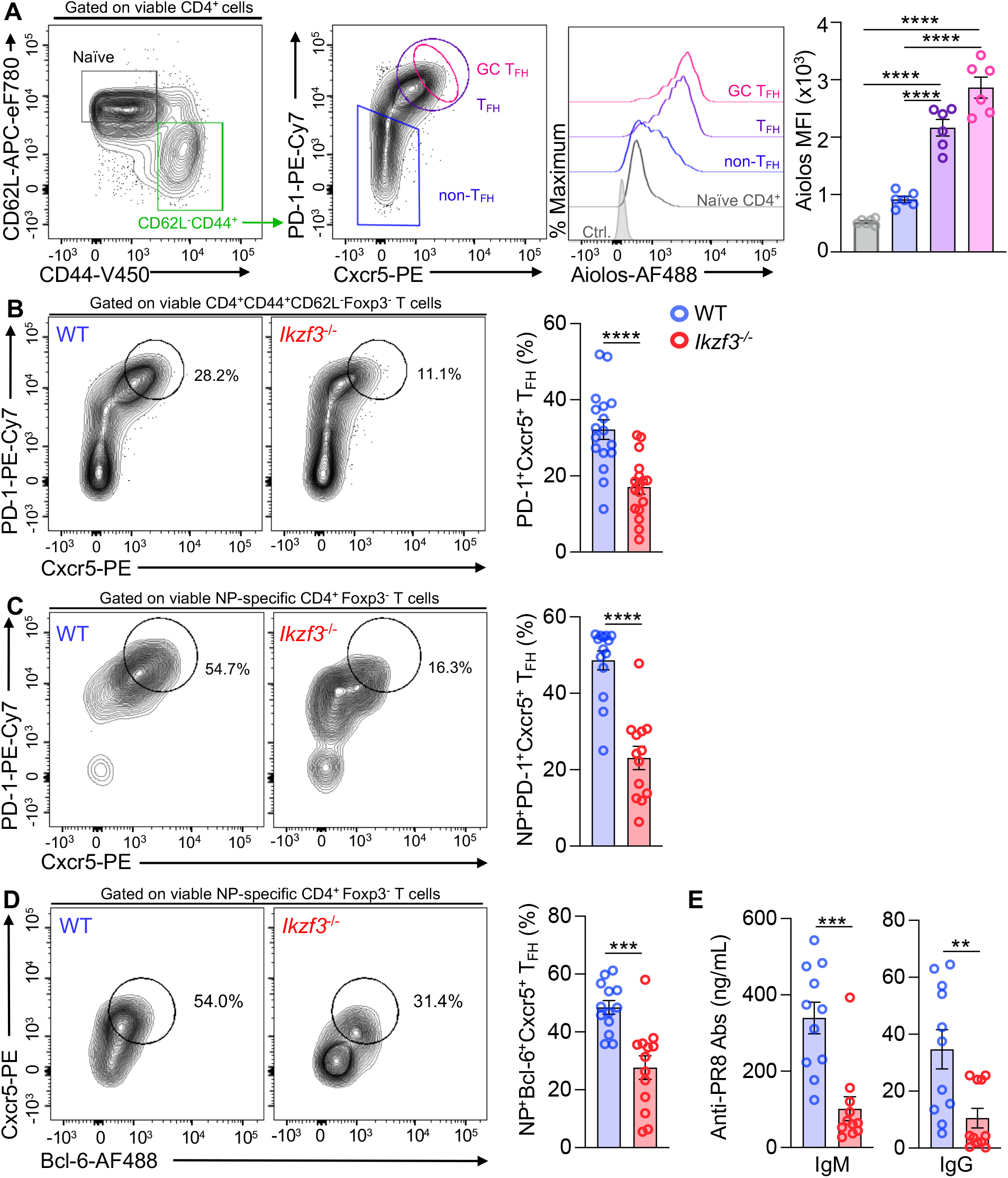
Aiolos deficiency results in disrupted T_FH_ cell differentiation and antibody production in response to influenza infection. (**A**) Naïve WT C57BL/6 mice were infected intranasally with 40 PFU influenza (A/PR8/34; “PR8”) for 8 days. Single-cell suspensions were generated from the draining (mediastinal) lymph node, and analysis of Aiolos protein expression in the indicated populations was performed via flow cytometry. Data are compiled from 2 independent experiments (n = 6 ± s.e.m; \*\*\*\**P* < 0.0001; unpaired Student’s t-test). (**B**) Analysis of the percentage of bulk PD-1^HI^Cxcr5^HI^ (T_FH_) populations generated during influenza infection in WT versus *Ikzf3^-/-^* mice. Data are compiled from 5 independent experiments (n = 17 ± s.e.m; \*\*\**P* < 0.001; unpaired Student’s t-test). (**C**) Analysis of the percentage of influenza nucleoprotein (NP)-specific PD-1^HI^Cxcr5^HI^ (T_FH_) populations generated in response to influenza infection. Following single-cell suspension, cells were stained with fluorochrome-labeled tetramers to identify NP-specific populations (n = 13-14 ± s.e.m; data are compiled from 4 independent experiments; \*\*\*\**P* < 0.0001; unpaired Student’s t-test). (**D**) Analysis of the percentage of influenza nucleoprotein (NP)-specific Bcl-6^HI^Cxcr5^HI^ (T_FH_) populations generated in response to influenza infection (n = 13 ± s.e.m; data are compiled from 4 independent experiments; \*\*\**P* < 0.001; unpaired Student’s t-test). (**E**) ELISA analysis of indicated serum antibody levels in ng/mL at 8 d.p.i. Data are compiled from 3 independent experiments (n = 11 ± s.e.m, \*\**P* < 0.01, ***< 0.001; unpaired Student’s t-test).

To determine whether Aiolos influences the generation of T_FH_ populations *in vivo*, we infected Aiolos-deficient (*Ikzf3^-/-^*) or WT mice with PR8 and assessed T_FH_ populations residing in the DLN via flow cytometry. Consistent with our hypothesis, *Ikzf3^-/-^* mice exhibited a significant decrease in the percentage of T_FH_ cells (identified as PD-1^+^Cxcr5^+^ or Bcl-6^+^Cxcr5^+^) relative to WT controls (**Fig. 1B** and **Extended Data Fig. 2A-C**). To determine whether the reduction of T_FH_ populations was maintained in antigen-specific populations, we stained with influenza nucleoprotein (NP311)-specific fluorochrome-labeled tetramers. As in bulk populations, loss of Aiolos resulted in a significant decrease in the percentage of NP-specific T_FH_ cells (**Fig. 1C,D)**. This reduction was not due to an overall difference in NP-specific cells in the absence of Aiolos, as these numbers were roughly equivalent between genotypes (**Extended Data Fig. 2D**). To determine whether the decrease in T_FH_ cell populations resulted in a functional impact on humoral responses, we measured influenza-specific antibody production via serum ELISA. We observed a significant reduction in flu-specific serum IgM and IgG in Aiolos-deficient animals relative to WT controls (**Fig. 1E**). Together, these findings are suggestive of a requirement for Aiolos in both the generation of antigen-specific T_FH_ cell populations and flu-specific antibody production during influenza virus infection.

### The T_FH_ transcriptional program is disrupted in the absence of Aiolos

We next evaluated Aiolos expression across *in vitro-generated* T helper cell subsets, including T_H_1, T_H_2, and TFH-like cells cultured in the presence of the murine TFH-inducing cytokine, IL-6^30,33,34^. Aiolos expression was significantly elevated in TFH-like cells compared to other cell types analyzed, and correlated with expression of Bcl-6 transcript and protein (**Fig. 2A** and **Extended Data Fig. 2B**). This was consistent with our previous findings^31^ and provided a robust *in vitro* system with which to determine mechanisms by which Aiolos supports the generation of T_FH_ cell populations. To examine alterations in the T_FH_ transcriptome in the absence of Aiolos, we cultured naive WT or Aiolos-deficient CD4^+^ T cells under T_FH_-polarizing conditions and performed RNA-seq analysis. Hierarchical clustering analysis revealed that Aiolos-deficient cells exhibited a distinct gene expression profile (**Extended Data Fig. 3A**). Genes downregulated in the absence of Aiolos included those encoding key pro-T_FH_ transcriptional regulators (*Bcl6, Tox, Tox2, Tcf7*, and *Zfp831*), as well as a number of other T_FH_-associated genes. Conversely, expression of the T_FH_ antagonist Blimp-1 was markedly increased in the absence of Aiolos (**Fig. 2B,C**). We further found that expression of genes encoding the IL-2 receptor subunits IL-2Rα (CD25) and IL-2Rβ (CD122) were elevated in Aiolos-deficient cells, while *Il6st* (encoding the IL-6R subunit gp130) was decreased (**Fig. 2B,D**). This was notable because IL-2/STAT5 signaling is a well-established repressor of T_FH_ differentiation and is known to induce expression of Blimp-1 ^11,12,26,29,35,36^. Consistent with these findings, gene set enrichment analysis (GSEA) of hallmark and immunologic signature gene sets revealed that genes upregulated in T_FH_ populations (versus T_H_1) were among the most downregulated in the absence of Aiolos, while those associated with the IL-2/STAT5 pathway were elevated (**Fig. 2E,F**).

**Figure 2.**
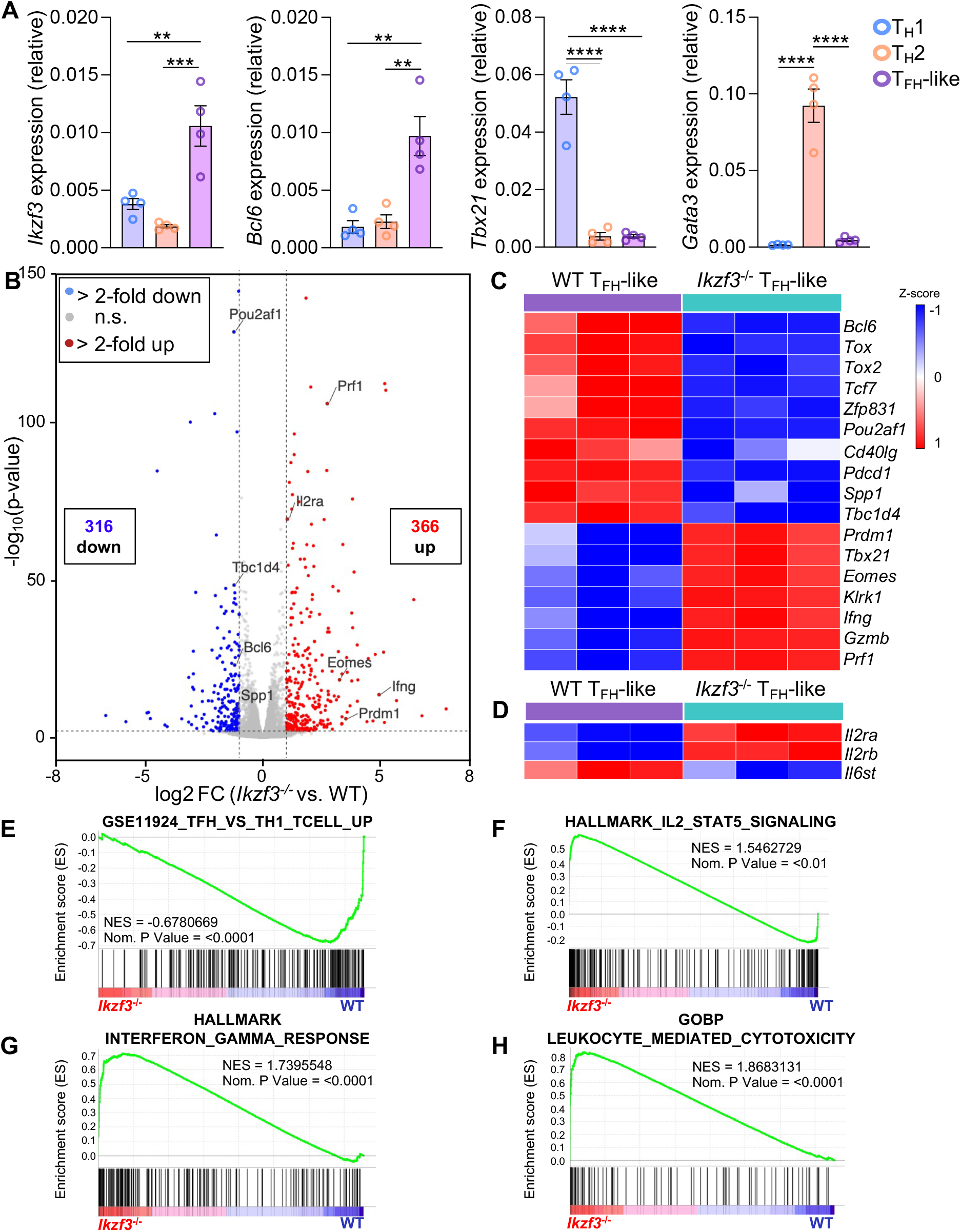
Loss of Aiolos results in alterations in the T_FH_ transcriptome. (**A**) Analysis of Aiolos transcript expression across T helper cell populations. Naïve murine CD4^+^ T cells were cultured under T_H_1, T_H_2, or TFH-polarizing conditions for 3 days. RNA was isolated and transcript analysis for the indicated genes was performed via qRT-PCR. Data are normalized and presented relative to the *Rps18* control. (n = 4 ± s.e.m; \*\**P* < 0.01, \*\*\**P* < 0.001, ****P < 0.0001; one-way ANOVA with Tukey’s multiple comparisons). (**B-H**) Naïve WT or Aiolos-deficient (*Ikzf3^-/-^*) CD4^+^ T cells were cultured under TFH-polarizing conditions for 3 days. RNA-seq analysis was performed to assess differentially expressed genes (DEGs) between WT and Aiolos-deficient cells. (**B**) Volcano plot displaying gene expression changes in Aiolos-deficient vs WT cells; genes of particular interest are labeled. Genes were color-coded as follows: no significant changes in expression (gray), upregulated genes with >2-fold change in expression with a P < 0.05 (red), downregulated genes with >2-fold change in expression with a P < 0.05 (blue) (**C**) Heat-map of DEGs positively and negatively associated with the T_FH_ gene program in WT vs. Aiolos-deficient cells; changes in gene expression are presented as relative expression by row (gene). (**E-H**) Pre-ranked (sign of fold change x −log_10_(p-value)) genes were analyzed using the Broad Institute Gene Set Enrichment Analysis (GSEA) software for comparison against ‘hallmark’, ‘gene ontology’, and ‘immunological signature’ gene sets. Enrichment plots for indicated gene sets are shown. Data are compiled from 3 biological replicates from 3 independent experiments.

Curiously, in contrast to the T_FH_ gene signature, we observed marked increases in expression of genes associated with T_H_1 and CD4-CTL differentiation and effector responses in the absence of Aiolos (**Fig. 2B,C**). These findings were supported by GSEA analysis indicating an increase in the expression of gene sets associated with both interferon gamma signaling and cytotoxic programming (**Fig. 2G,H**). Collectively, these findings suggested that Aiolos may not only promote T_FH_ differentiation through activation of key transcriptional regulators, but it may also safeguard the T_FH_ gene program by repressing genes associated with alternative (T_H_1, CD4-CTL) cell states.

### Aiolos associates with and alters chromatin structure at T_FH_ gene regulatory elements

RNA-seq and confirmatory qRT-PCR analyses revealed reduced expression of several key T_FH_ transcriptional regulators in Aiolos-deficient cells cultured under T_FH_-like polarizing conditions (**Fig. 2C** and **3A-C**). To determine whether Aiolos directly regulated these gene targets, we utilized a combined Assay for Transposase-Accessible Chromatin (ATAC)-seq and chromatin immunoprecipitation (ChIP) approach. We began by leveraging ATAC-seq analysis of *in vitro* polarized WT and *Ikzf3^-/-^* CD4^+^ T cells to identify differing regions of chromatin accessibility (**Fig. 3D-F**). We found several regions in which chromatin accessibility was negatively impacted by the absence of Aiolos, including those at the *Bcl6* promoter, *Tcf7* 5’UTR, and a region upstream of the *Zfp831* TSS. Bcl-6 and TCF-1 (Tcf7) are well-established positive regulators of T_FH_ cell differentiation, while Zfp831 was recently identified as a direct inducer of both of these factors^11,16,17,19,21–23^. To determine whether Aiolos may directly regulate these genes in *in vitro-* generated TFH-like cells, we performed ChIP analyses at identified differentially accessible regions and control regions which did not display altered chromatin accessibility. Consistent with our hypothesis, Aiolos enrichment was elevated at *Bcl6, Tcf7*, and *Zfp831* regulatory regions relative to that of an irrelevant antibody control and that observed at control genomic regions (**Fig. 3G-I**).

**Figure 3.**
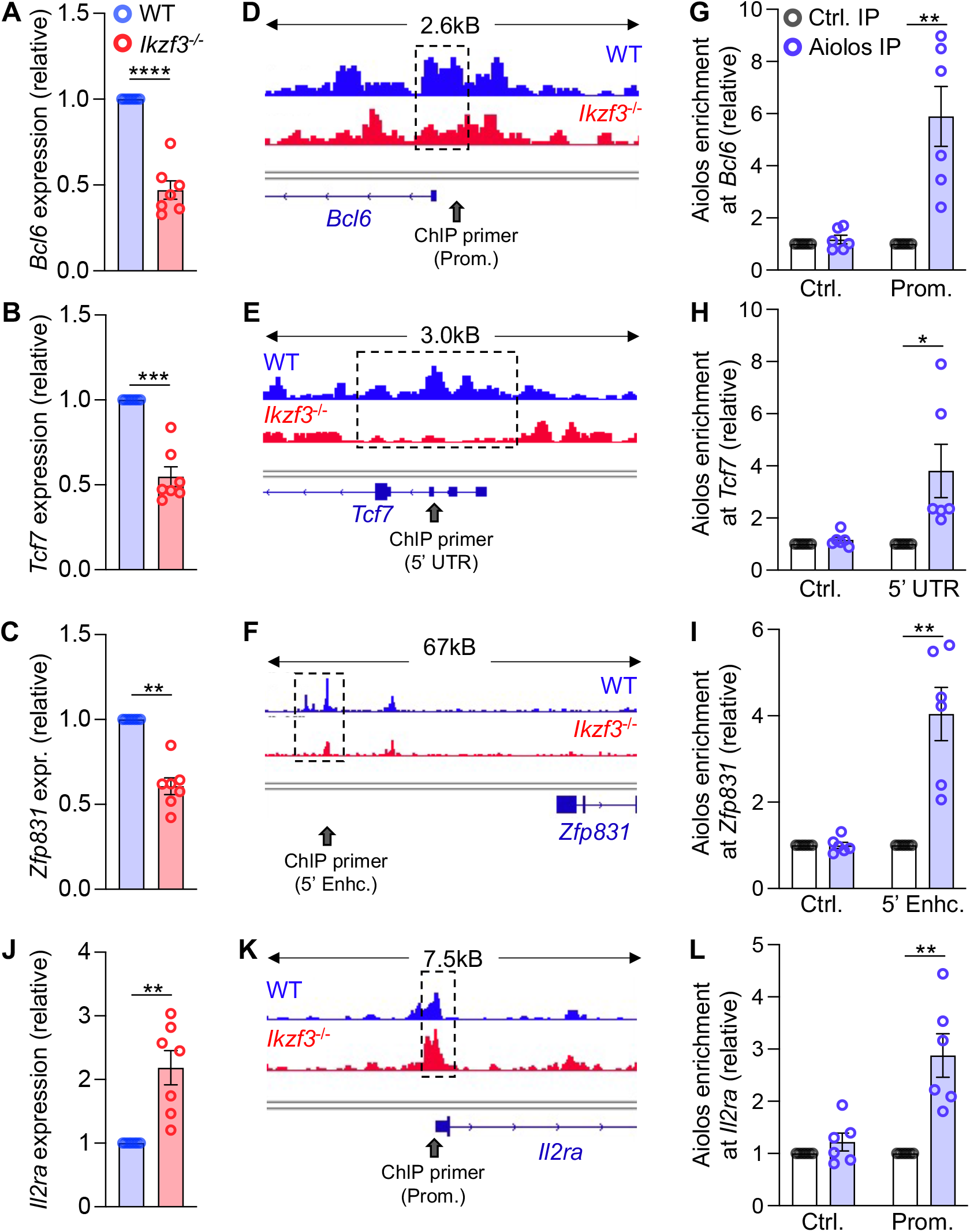
Aiolos is enriched at key T_FH_ transcription factor gene loci. **(A-C, J)** Naïve CD4^+^ T cells were cultured under TFH-like polarizing conditions. After 3 days, transcript analysis for the indicated genes was performed via qRT-PCR. (n=7 ± s.e.m; \*\**P* < 0.01, ***<0.001, ****< 0.0001; unpaired Student’s t-test). **(D-F, K)** Assay for Transposase-Accessible Chromatin (ATAC)-seq was performed on WT and Aiolos-deficient CD4^+^ cells cultured under TH1-skewing conditions to assess changes in accessibility at differentially expressed gene loci. CPM-normalized representative images from two independent experiments are shown. Scales are identical between WT and Aiolos-deficient tracks for each locus analyzed (range = 0.5-2.0 CPM). Approximate locations of primers used for chromatin immunoprecipitation (ChIP) are indicated with gray arrows. **(G-I, L)** ChIP analysis of Aiolos enrichment at the indicated regulatory regions, or negative control regions, are shown. Data were normalized to total IgG and presented relative to enrichment in the IgG IP control sample. (n=6 ± s.e.m; \**P* < 0.05, \*\**P* < 0.01; unpaired Student’s t-test).

Our RNA-seq and qRT-PCR analyses further revealed that TFH-skewed Aiolos-deficient cells express significantly altered cytokine receptor repertoires, including increased expression of *Il2ra* (**Figs. 2D** and **3J**). Strikingly, ATAC-seq analysis revealed increased chromatin accessibility at the *Il2ra* promoter in *Ikzf3^-/-^* compared to WT cells, while accessibility was decreased at T_FH_ transcription factor loci (**Fig. 3D-F,K**). As Aiolos has also been implicated in the repression of target genes, we hypothesized that Aiolos may also function to repress *Il2ra* expression. Indeed, ChIP analysis revealed that Aiolos was enriched at the *Il2ra* promoter relative to an upstream control region (**Fig. 3L**). Given that Blimp-1 (*Prdm1*) expression was also increased in the absence of Aiolos, we considered the possibility that Aiolos may directly regulate its expression. While a known *Prdm1* intronic enhancer element exhibited increased accessibility in the absence of Aiolos, we did not observe Aiolos enrichment at this region (**Extended Data Fig. 4A,B**). This suggests that Aiolos may repress Blimp-1 expression via an indirect mechanism or possibly through interaction with another regulatory region (**Extended Data Fig. 4C**). Taken together, these data are suggestive of a mechanism whereby Aiolos directly regulates key T_FH_ transcription factors and CD25 expression in a reciprocal fashion to promote T_FH_ differentiation.

### Aiolos deficient cells express hallmark CD4-CTL transcription factors and molecules

RNA-seq analyses of Aiolos-deficient cells cultured under TFH-like polarizing conditions revealed not only compromised T_FH_ cell programming, but also augmented expression of transcription factors and effector features associated with T_H_1 and CD4-CTLs (**Fig. 2B,C**). These data are consistent with other studies demonstrating that regulators of T_FH_ differentiation oppose both T_H_1 and CD4-CTL differentiation^15,17^. We first sought to examine the role of Aiolos in regulating T_H_1 differentiation by culturing naive WT and Aiolos-deficient CD4^+^ cells under T_H_1 cell polarizing conditions. We assessed expression of *Bcl6* and the T_H_1 lineage-defining transcription factor T-bet (encoded by *Tbx21*) at the transcript and protein level. Although we observed diminished expression of *Bcl6* in the absence of Aiolos, *Tbx21* transcript and protein remained unchanged, suggesting that loss of Aiolos may not augment T_H_1 differentiation in this context (**Fig. 4A,B**). Strikingly, however, expression of the related T-box factor, Eomesodermin (Eomes), which has a stronger association with the cytotoxic programs of CD4^+^ and CD8^+^ T cells, was significantly increased (**Fig. 4A,B**)^4,7,37,38^.

**Figure 4.**
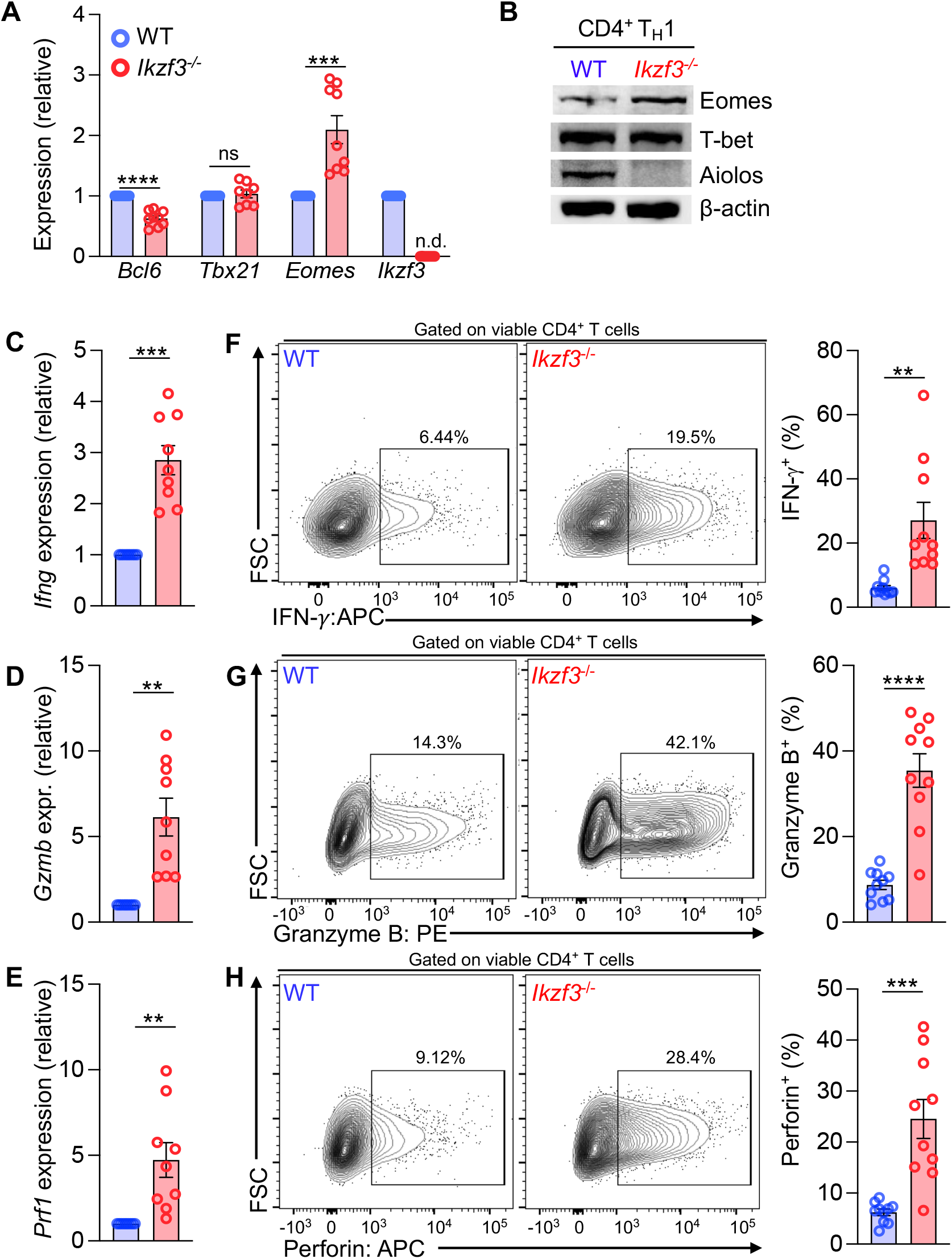
Expression of the CD4-CTL gene program and effector molecule production are increased in the absence of Aiolos. Naive WT or Aiolos-deficient (*Ikzf3^-/-^*) CD4^+^ T cells were cultured under TH1-polarizing conditions for 3 days. (**A, C-E)** Transcript analysis for the indicated genes was performed via qRT-PCR. Data are normalized to *Rps18* control and presented relative to the WT sample for each gene. (n=9 ± s.e.m; **P < 0.01, ***P < 0.001, ****P < 0.0001; unpaired Student’s t-test; n.d., not detected). **(B)** Immunoblot analysis of the indicated proteins. β-actin serves as a control to ensure equivalent protein loading. A representative image from 3 independent experiments is shown. **(F-H)** Analysis of cytotoxic effector molecule production by flow cytometry. Cells were stimulated for 3h with PMA and Ionomycin in the presence of protein transport inhibitors. Cells were fixed and permeabilized and stained for the indicated proteins. (n=10 ± s.e.m; **P < 0.01, ***P < 0.001, ****P < 0.0001; unpaired Student’s t-test).

Relative to T_H_1 populations, CD4-CTLs display increased production of IFN-γ, as well as the expression of cytolytic molecules including granzymes and perforin^4^. Consistent with increased Eomes expression, we found that *Ikzf3^-/-^* cells polarized under T_H_1 conditions expressed more IFN-γ, granzyme B, and perforin at both the transcript and protein levels (**Figs. 4C-H**). To better understand transcriptomic changes, we performed RNA-seq analysis of WT and Aiolos-deficient cells cultured under T_H_1 conditions. In agreement with the prior data, this analysis revealed increased expression of additional transcription factors (*Prdm1, Runx3*) and cell surface receptors associated with the cytotoxic programs of CD4^+^ T cells (**Fig. 5A-C**). In contrast, transcription factors associated with T_FH_ differentiation and antagonists of CTL-differentiation were downregulated in the absence of Aiolos (**Fig. 5C**). Further, gene set enrichment analysis revealed that gene signatures associated with cytotoxicity and IL-2/STAT5 signaling were among the most upregulated signatures in the absence of Aiolos (**Fig. 5D,E**). Collectively, these data indicate that Aiolos deficiency results in an increase in hallmark features of cytotoxicity and supports a role for Aiolos as a repressor of the cytotoxic gene program in CD4^+^ T cells.

**Figure 5.**
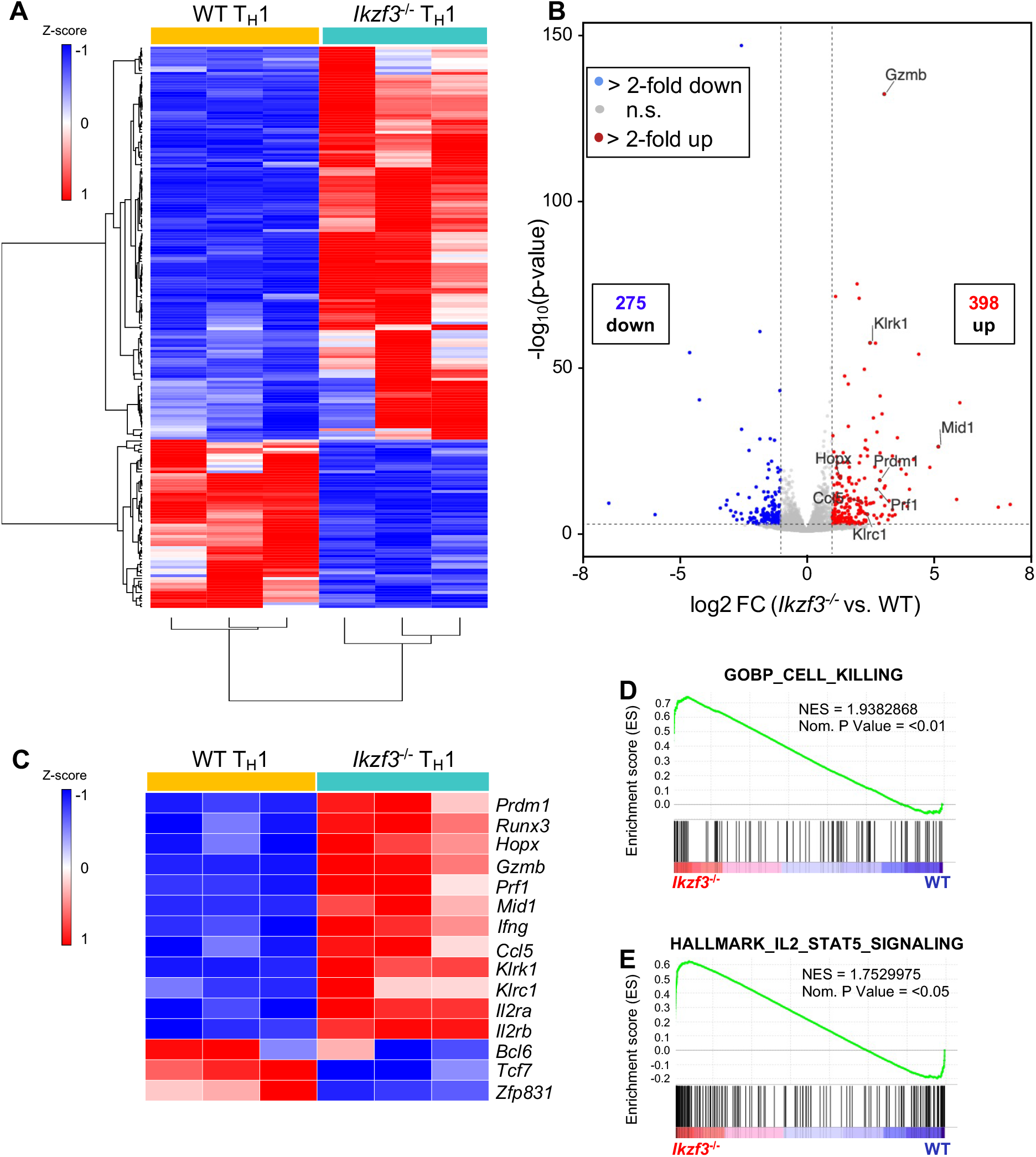
Aiolos deficiency results in alterations in CD4^+^ cytotoxic programming *in vitro.* Naïve WT or Aiolos-deficient (*Ikzf3^-/-^*) CD4^+^ T cells were cultured under TH1-polarizing conditions for 3 days. RNA-seq analysis was performed to assess differentially expressed genes (DEGs) between WT and Aiolos-deficient cells. (**A**) Heatmap of the top 200 most differentially expressed genes (p < 0.05). Genes and samples were clustered using Euclidean distance. Samples are compiled from 3 biological replicates from 3 independent experiments. (**B**) Volcano plot displaying gene expression changes in Aiolos-deficient vs WT cells; genes of particular interest are labeled. Genes were color-coded as follows: no significant changes in expression (gray), upregulated genes with >2-fold change in expression with a P < 0.05 (red), downregulated genes with >2-fold change in expression with a P < 0.05 (blue). (**C**) Heatmap of DEGs positively and negatively associated with the T_H_1 gene program in WT vs. Aiolos-deficient cells; changes in gene expression are presented as relative expression by row (gene). (**D-E**) Pre-ranked genes were analyzed using the Broad Institute Gene Set Enrichment Analysis (GSEA) software for comparison against ‘hallmark’ and ‘gene ontology’ gene sets. Enrichment plots for indicated gene sets are shown. Data are compiled from 3 biological replicates from 3 independent experiments.

### Aiolos deficiency results in acquisition of cytotoxic features by CD4 T cells *in vivo*

Our prior *in vivo* data revealed a reduction in the percentage of T_FH_ cells in the DLN of Aiolos-deficient mice, while our *in vitro* studies demonstrated an increase in cytotoxic programming in the *Ikzf3^-/-^* cells cultured under T_H_1 or T_FH_-like conditions. To further determine the impact of Aiolos-deficiency *in vivo*, we next infected WT and Aiolos-deficient (*Ikzf3^-/-^*) mice with PR8 and assessed T-bet and Eomes expression in NP-specific CD4^+^ T cells of the DLN via flow cytometry. Consistent with our *in vitro* findings, there was no difference in the percentage of T-bet positive NP-specific cells from the DLN of WT and *Ikzf3^-/-^* mice (**Figs. 6A,B**). However, the percentage of Eomes positive NP-specific cells was enhanced in the DLN of Aiolos-deficient mice compared to that of WT (**Figs. 6A,B**). We next examined perforin expression, an Eomes target gene and CTL effector molecule, and found elevated percentages of perforin producing Eomes^+^T-bet^+^ and Eomes^+^T-bet^-^ cells in Aiolos-deficient mice compared to WT (**Fig. 6C,D**). In contrast, we did not observe a difference in Eomes^-^T-bet^+^ perforin expressing cells, demonstrating a correlation between Aiolos-deficiency and Eomes and perforin expression (**Fig. 6D**). Similarly, analysis of NP-specific populations revealed increased percentages of Eomes^+^ perforin expressing cells in *Ikzf3^-/-^* mice compared to WT (**Fig. 6E**). We also examined expression of the CTL marker Cx3cr1 and found increased percentages of NP-specific Cx3cr1 ^+^ cells in Aiolos-deficient mice relative to WT (**Fig. 6F** and **Extended Data Fig. 5**). Collectively, these *in vitro* and *in vivo* findings indicate that Aiolos restrains expression of a cytotoxic-like program, at least in part, via repressive effects on the T-box transcription factor Eomes.

**Figure 6:**
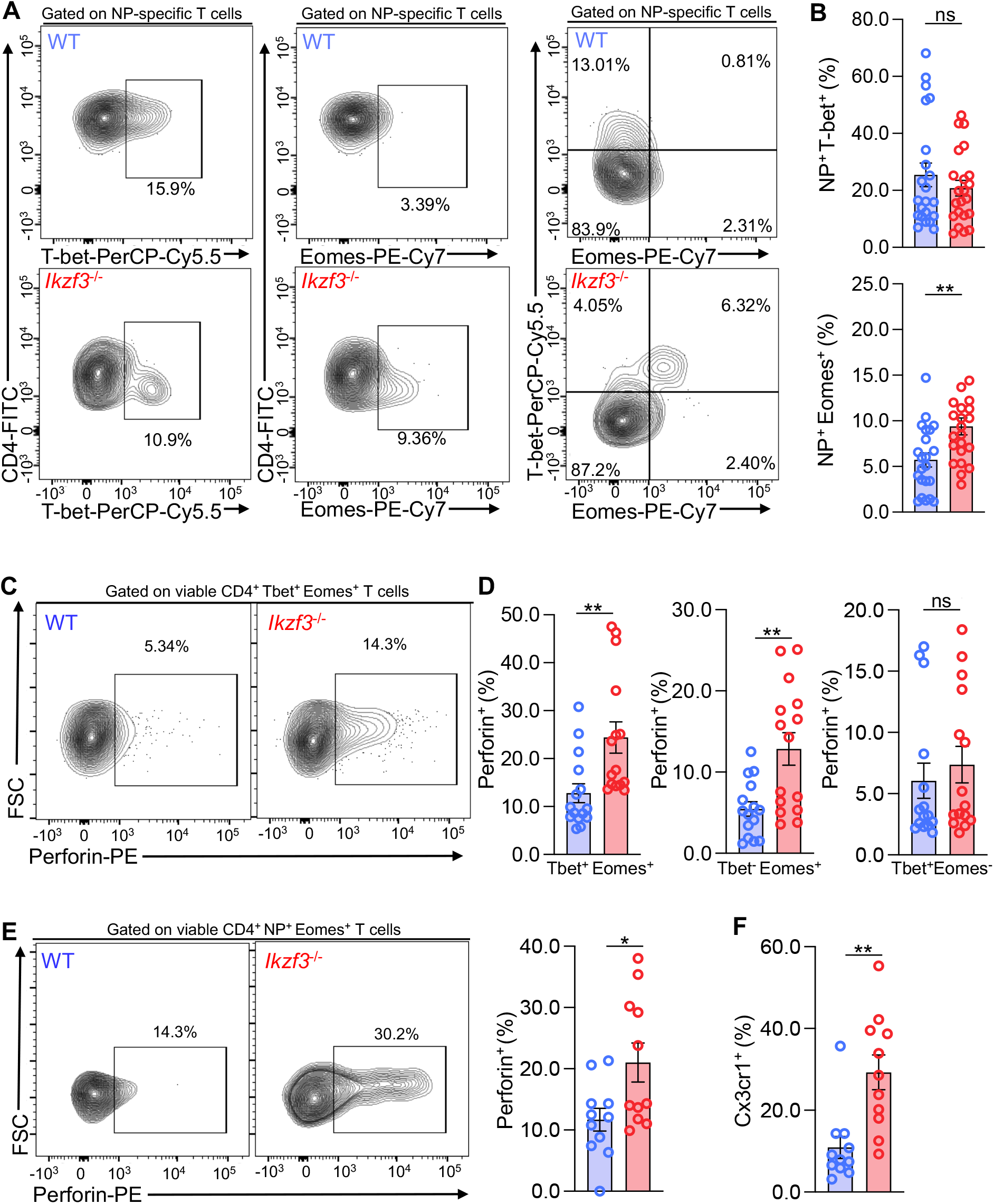
Aiolos deficiency results in acquisition of cytotoxic features by CD4^+^ T cells *in vivo.* Naïve WT or Aiolos-deficient (*Ikzf3^-/-^*) mice were infected intranasally with 40 PFU influenza (A/PR8/34; “PR8”). After 8 days, draining (mediastinal) lymph nodes were harvested and viable CD4^+^ effector populations were analyzed via flow cytometry. (**A-B**) Analysis of influenza nucleoprotein (NP)-specific cells for expression of T-bet and Eomes. Data are compiled from 7 independent experiments (n=22 ± s.e.m; **P < 0.01; unpaired Student’s t-test). (**C-E**) Single-cell suspensions from the DLN were incubated in culture medium in the presence of protein transport inhibitors for 2-3 hours. Perforin production by T-bet^+^Eomes^+^ and Eomes^+^ populations was analyzed via flow cytometry. Data are compiled from 4 independent experiments. (n = 14 ± s.e.m; *P < 0.05, **P < 0.01; unpaired Student’s t-test). (**F**) Analysis of Cx3cr1 surface expression on NP^+^Eomes^+^cells. Data are compiled from 4 independent experiments (n = 14 ± s.e.m; **P < 0.01; unpaired Student’s t-test).

### Loss of Aiolos results in elevated CD25 expression

IL-2/STAT5 signaling is known to both repress T_FH_ and promote CD4-CTL differentiation. Since our RNA-seq analysis indicated that IL-2 receptor subunit transcript was elevated in the absence of Aiolos, we next examined whether Aiolos may both support T_FH_ generation and repress CD4-CTL differentiation by repressing aspects of IL-2/STAT5 signaling *in vivo*. We infected Aiolos-deficient mice with PR8 and assessed alterations in the surface expression of CD25 on CD4^+^ effector T cells (CD44^+^CD62L^-^Foxp3^-^) of the DLN. Indeed, analysis of bulk CD4^+^ effector populations revealed a significant increase in both the percentage of CD25^+^ cells and the level of CD25 expression in the absence of Aiolos (**Fig. 7A**). Analysis of antigen-specific effector cells from the DLN revealed a similar upregulation of CD25 on *Ikzf3^-/-^* versus WT cells (**Fig. 7B**). We considered the possibility that the increase in CD25 surface expression may be due to an overall increase in IL-2 production in the absence of Aiolos, as Aiolos was previously shown to repress IL-2 production in *in vitro*-generated T_H_17 cells^39^. However, analysis of IL-2 production *in vivo* revealed no significant difference between WT and Aiolos-deficient CD4^+^ T cell populations (**Fig. 7C**). These data, together with our identification of *Il2ra* as a direct Aiolos target, suggest that Aiolos represses IL-2 responsiveness to both promote T_FH_ and repress CD4-CTL differentiation.

**Figure 7.**
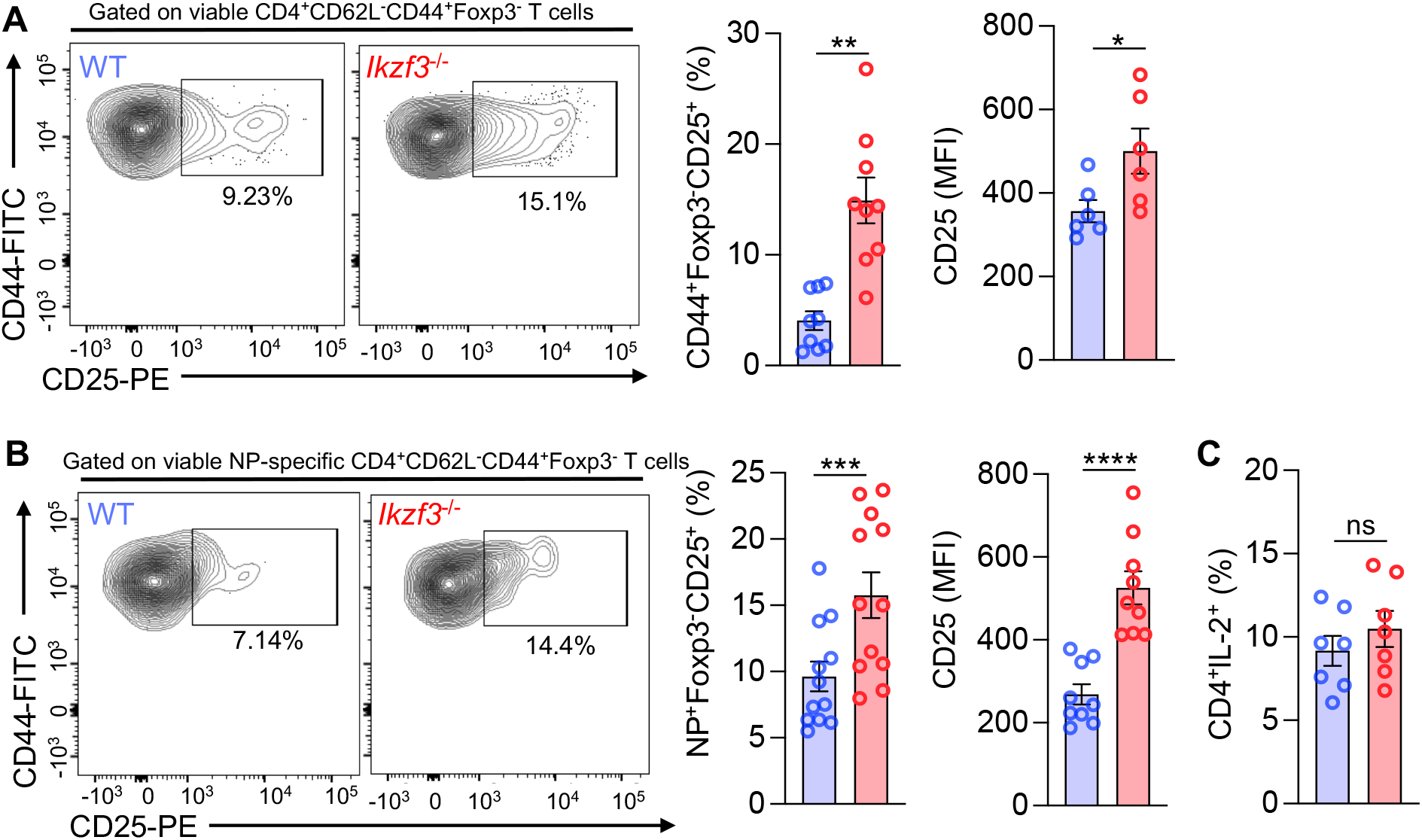
Loss of Aiolos results in elevated expression of CD25. Naïve WT or Aiolos-deficient (*Ikzf3^-/-^*) mice were infected with influenza (PR8) for 8 days. (**A-B**) CD25 (IL-2Rα) surface expression was evaluated on viable CD4^+^CD44^+^CD62L^-^Foxp3^-^ (effector) T cells from the DLN via flow cytometry. Percentage of CD25^+^ cells and median fluorescence intensity (MFI) are shown for bulk (A) and antigen-specific (B) cell populations. Data are compiled from 3 independent experiments. (n = 9 ± s.e.m; *P < 0.05, **P < 0.01, ***P < 0.001, ****P <0.0001; unpaired Student’s t-test). (**C**) Production of IL-2 by CD4^+^ T cell populations was analyzed in WT vs. Aiolos-deficient mice by flow cytometry. Data are compiled from 2 independent experiments. (n = 7 ± s.e.m; unpaired Student’s t-test).

### Absence of Aiolos results in increased STAT5 enrichment at CTL genes

IL-2/STAT5 signaling is known to positively regulate the cytotoxic gene program in CD4^+^ T cells. As such, we considered the possibility that increased CD25 expression in the absence of Aiolos may lead to increased STAT5 activation and STAT5-dependent induction of the CTL gene program. Analysis of tyrosine phosphorylation of STAT5 revealed increased STAT5 activation in Aiolos-deficient compared to WT cells cultured under T_H_1 conditions (**Fig. 8A**). ATAC-seq analysis of the known STAT5 target genes, *Il2ra* and *Prdm1*, revealed increased chromatin accessibility at promoter and intronic enhancer elements in the absence of Aiolos, respectively, and these alterations positively correlated with increased gene expression (**Fig. 8B,C** and **Extended Data Fig. 6A,B**). The areas of increased accessibility also corresponded to known sites of STAT5 enrichment, as determined by previously published ChIP-seq data (**Fig. 8B,C**). To determine whether STAT5 enrichment was enhanced at these regions in the absence of Aiolos, we performed ChIP analysis in WT and *Ikzf3^-/-^* cells. Indeed, ChIP revealed enhanced STAT5 association with both regulatory elements in the absence of Aiolos (**Fig. 8D,E**). Similar analyses revealed enhanced chromatin accessibility and STAT5 enrichment at regulatory regions of *Eomes, Ifng, Gzmb*, and *Prf1* in Aiolos-deficient cells (**Fig. 8F,G** and **Extended Data Fig. 6C**). Collectively, the above findings support a mechanism whereby Aiolos deficiency results in augmented IL-2Rα expression, enhanced sensitivity to IL-2, and increased association of STAT5 with CD4-CTL gene targets.

**Figure 8.**
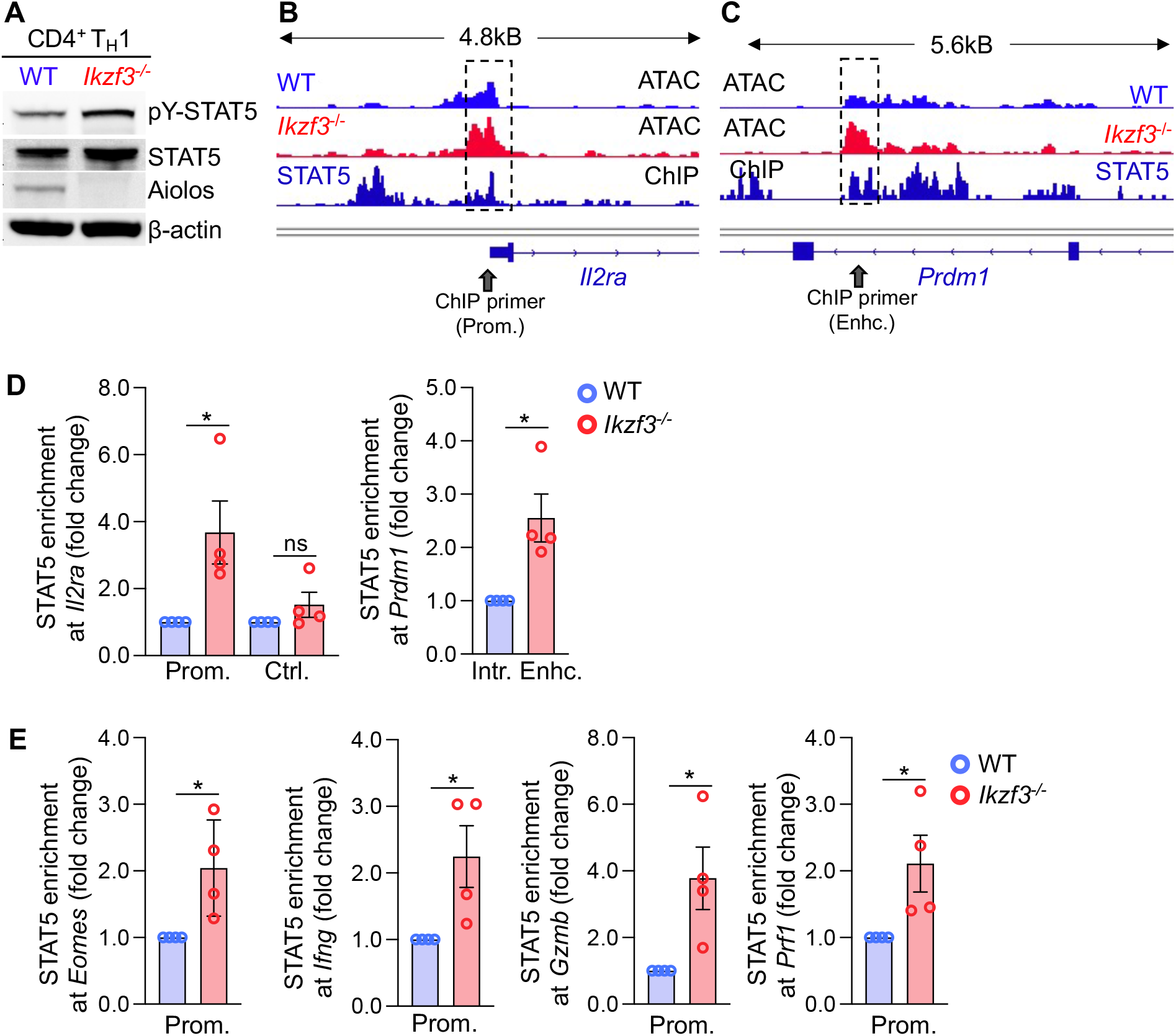
Aiolos deficiency results in alterations in chromatin accessibility and STAT5 binding at regulatory regions of CD4-CTL target genes. Naïve WT and Aiolos-deficient CD4^+^ T cells were cultured in the presence of T_H_1 polarizing conditions for 3 days. **(A)** Immunoblot analysis of the indicated proteins. β-actin serves as a loading control. A representative image from 3 independent experiments is shown. **(B-C)** ATAC-seq analyses of T_H_1 samples overlaid with published STAT5 ChIP-seq data. WT (light blue, top track) and Aiolos-deficient (red, middle track) samples are displayed as CPM-normalized Integrative Genomics Viewer (IGV) tracks (representative from 2 independent experiments). STAT5 ChIP-seq data alignment (dark blue) is shown in the bottom track (from GEO # GSE102317 (GSM2734684)). Regulatory regions of note are indicated by dashed boxes. Approximate locations of primers used to assess STAT5 enrichment by ChIP are indicated with gray arrows. (**D-E**) Naïve WT and Aiolos deficient CD4^+^ T cells were cultured in the presence of T_H_1 polarizing conditions for 4 days. ChIP analysis was performed for STAT5 and an IgG control at the indicated regions. Data are normalized to total input sample. IgG values were subtracted from percent enrichment and data are displayed relative to the WT sample. (n = 4 ± s.e.m; *\*P* <0.05; unpaired Student’s t-test).

## Discussion

Our present study identifies Aiolos as a reciprocal regulator of the differentiation programs of T_FH_ and CD4-CTL cells. Mechanistically, we find that Aiolos associates with and positively regulates chromatin structure at the regulatory elements of genes encoding Bcl-6, TCF-1, and Zfp831, transcription factors required for T_FH_ differentiation. We further show that Aiolos represses CD25 expression and thus cell responsiveness to IL-2, a potent T_FH_ inhibitory cytokine. Consequently, we find that Aiolos-deficiency results in increased IL-2/STAT5 signaling, repression of T_FH_ differentiation, and induction of the cytotoxic gene program. Importantly, a key regulatory aspect of the increased cytotoxic programming of these cells is induction of Eomes expression via IL-2/STAT5 signaling, rather than impacts on the related T-box factor, T-bet.

The findings presented here indicate that Aiolos promotes T_FH_ differentiation by at least two mechanisms. First, Aiolos directly associates with the regulatory regions of key T_FH_ transcriptional regulators. Since the identification of Bcl-6 as the lineage-defining factor for T_FH_ differentiation over a decade ago, there has been steady growth in our understanding of the transcriptional mechanisms that regulate T_FH_ differentiation^11,16,17^. This includes the identification of additional transcription factors that operate upstream or in parallel with Bcl-6, such as TCF-1 and Zfp831, to promote expression of the T_FH_ gene program^19,21–23^. Our findings demonstrate that Aiolos is enriched at all three genes and that expression of these genes, as well as T_FH_ differentiation, is disrupted in its absence. Importantly, we also find that expression of the T_FH_ antagonist, Blimp-1, is enhanced in the absence of Aiolos. Taken together, these findings indicate that Aiolos represents a key upstream transcriptional regulator of other transcription factors that both promote and repress T_FH_ differentiation.

Second, our transcriptomics data indicate that Aiolos is a reciprocal regulator of subunits of the IL-6R and IL-2R complexes, which allow for the propagation of cytokine signaling pathways that are known to promote and repress T_FH_ differentiation, respectively^14^. These data suggest that Aiolos positively impacts T_FH_ differentiation by maintaining a pro-TFH balance of these cytokine receptors. Further highlighting the relationship between Aiolos and IL-6R signaling are prior findings from our lab that identified a novel transcriptional complex composed of Aiolos and STAT3 which, among other cytokines, is activated in response to IL-6^31^. Thus, our combined findings suggest that Aiolos regulates not only IL-6R expression, but perhaps also the downstream activities of STAT3 in the nucleus. From this standpoint, it is interesting to speculate whether Aiolos/STAT3 complexes may regulate gene expression by antagonizing STAT5 association with target genes. Indeed, prior studies have demonstrated that competitive mechanisms between STAT3 and STAT5 are important in the promotion and inhibition of T_H_17 and T_FH_ differentiation^12,26,29,30,40,41^. To this end, it is notable that we find Aiolos enrichment at regulatory regions known to be targeted by STAT5. This includes the promoters of *Bcl6* and *Il2ra*, genes whose expression we show are inversely regulated by the activities of Aiolos and STAT5. Further work is needed to elucidate the mechanistic interplay between these transcription factors throughout the genomes of the cells studied here, as well as other IL-6 and IL-2 responsive immune cell populations.

It is also interesting to compare the known role of Aiolos in the differentiation of T_H_17 cells to that of our findings with T_FH_ populations. In a prior study, Aiolos was found to be a direct repressor of IL-2 expression^39^. This ultimately aided in the differentiation of T_H_17 cells due to a lessening of the negative impacts of IL-2 on T_H_17 cell formation. Alternatively, we find here that Aiolos represses CD25 expression, thus limiting the ability of differentiating T_FH_ cells to respond to signals from this cytokine. Our findings are consistent with those from other studies demonstrating that T_FH_ cells function as IL-2 producers during responses to infection and that T_FH_ cells rely on signals from IL-6/STAT3 to allow for repression of IL-2R subunits^30,33,42^.

Our findings are also significant as they pinpoint Aiolos as a repressor of the cytotoxic gene program in CD4^+^ T cells. CD4-CTLs are diverse regulators of immune responses as they are known contributors during infection, anti-tumor immunity, and autoimmunity^4^. To date, the mechanisms that promote CD4-CTL differentiation have remained somewhat enigmatic with the apparent formation of these cells driven via multiple signaling pathways and sets of transcriptional regulators^3,4^. Our current finding that Aiolos drives Bcl-6 and TCF-1 expression is consistent with a prior study demonstrating that these factors can repress the acquisition of cytotoxic features by CD4^+^ T cells^15^. Prior work has also demonstrated that IL-2/STAT5 signaling and increased Blimp-1 expression can help to drive cytotoxic programming in CD4^+^ T cells^13,43,44^. Indeed, we find that increased IL-2 responsiveness in the absence of Aiolos plays a central role by enhancing the association of STAT5 with key cytotoxic gene regulatory elements. Importantly, we also find that increased STAT5 association correlates with enhanced chromatin accessibility at these genes. Notably, this mechanism appears to drive the expression of the genes encoding both Eomes and Blimp-1, known positive regulators of the cytotoxic program in CD4^+^ T cells. We further find enhanced STAT5 association with genes encoding effector cytokines and molecules including IFN-γ, granzyme B, and perforin. Interestingly, alterations to the cytotoxic gene program appear to be largely driven by IL-2/STAT5-dependent increases in Eomes expression, rather than impacting T-bet expression. In this regard, it will be important in future studies to explore the relationship between Aiolos and both T-box factors in other immune cellular settings, such as CD8^+^ T cell populations where the balance of these transcription factors is critical in promoting diverse effector responses, exhaustion, and memory cell programming^45–48^.

Finally, it should be noted that Aiolos is currently a target of the FDA approved chemotherapeutic lenalidomide for the treatment of multiple myeloma^49^. Further work has shown that exposure of CAR-T cells to lenalidomide can potentiate anti-tumor activity of these cells in murine cancer models, though the mechanisms are still being investigated^50^. Still, the effect of lenalidomide on cells across the immune system and the mechanisms involved are currently unclear. Our findings identifying Aiolos as a pivotal regulator of T_FH_ and CD4-CTL differentiation provide needed clarity into the impact of Aiolos on immune cell populations and highlight its potential as a therapeutic target for the manipulation of CD4^+^ T cell-dependent humoral and cytotoxic responses.

## Materials and Methods

### Mouse strains

C57BL/6 mice were obtained from the Jackson Laboratory. *Ikzf3^-/-^* mice were originally obtained from Riken BRC and backcrossed to the C57BL/6 Jackson background for at least 10 generations to generate *Ikzf3*^-/-^/J mice.

### CD4^+^ T cell isolation and culture

Naive CD4^+^ T cells were isolated from the spleens and lymph nodes of 5-7 week old mice using the BioLegend Mojosort naïve CD4^+^ T cell isolation kit according to the manufacturer’s recommendations. Naïve cell purity was verified by flow cytometry and routinely exceeded 96-98%. For the *in vitro* polarization of TFH-like and T_H_1 populations, cells (1.5-2×10^5^ cells/mL) were cultured in complete IMDM ((IMDM [Life Technologies], 10% FBS [26140079, Life Technologies], 1% Penicillin-Streptomycin [Life Technologies], and 50mM 2-mercapto-ethanol [Sigma-Aldrich]) on plate-bound anti-CD3 (clone 145-2C11; 5ug/mL) and anti-CD28 (clone 37.51; 2ug/mL) in the presence of IL-4 neutralizing antibody (11B11, BioLegend, 5ug/mL) for 18-20 hours before the addition of cytokines as follows: for T_FH_ polarization: rmIL-6 (R&D, 100ng/mL); for T_H_1 polarization, rmIL-12 (R&D, 5ng/mL) and rhIL-2 (NIH, 50U/mL). For T_H_2 polarization, cells were cultured on plate-bound anti-CD3 (clone 145-2C11; 5ug/mL) and anti-CD28 (clone 37.51; 2ug/mL) in the presence of IFN-γ neutralizing antibody (XMG1.2, BioLegend, 10ug/mL) and rmIL-4 (R&D, 10ng/mL). For some experiments, IL-2 was also neutralized (JES6-1A12, BioLegend, 5ug/mL). Cells were cultured on stimulation for 72-96h before analysis.

### RNA isolation and qRT-PCR

Total RNA was isolated from the indicated cell populations using the Macherey-Nagel Nucleospin RNA Isolation kit as recommended by the manufacturer. cDNA was generated from mRNA template using the Superscript IV First Strand Synthesis System with provided oligo dT primer (Thermo Fisher). qRT-PCR reactions were performed with the SYBR Select Mastermix for CFX (ThermoFisher) using 10-20ng cDNA per reaction and the primers provided below. All qRT-PCR was performed on the CFX Connect (BioRad). Data were normalized to *Rps18* and presented either relative to *Rps18* or relative to the control sample, as indicated.

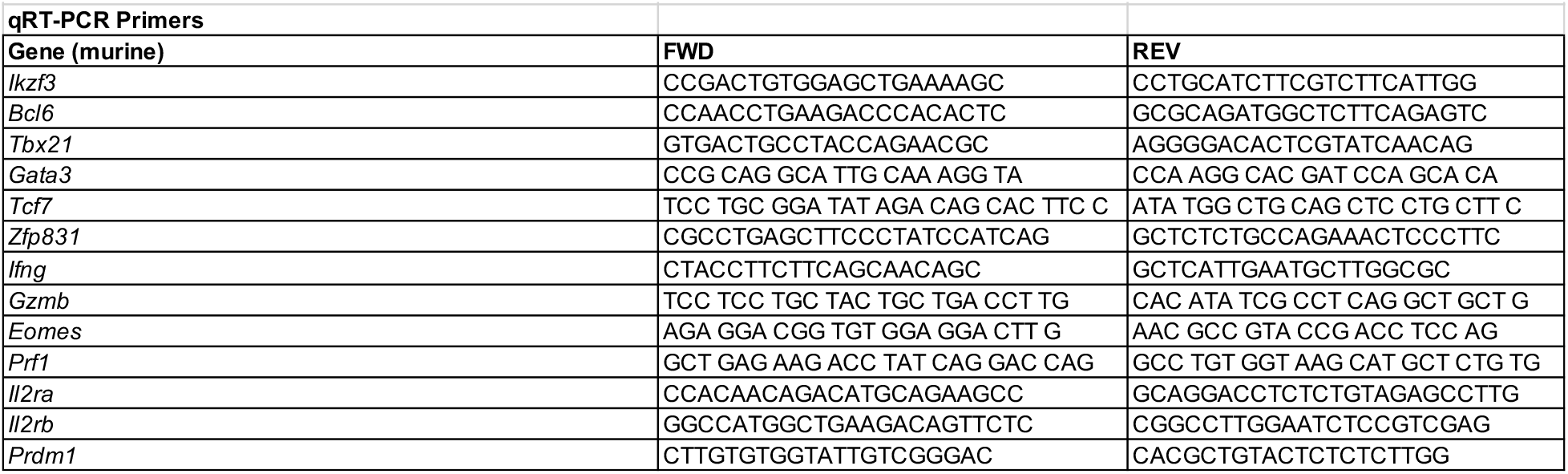

### Immunoblot analysis

Immunoblots were performed as described previously^31,51^. Briefly, cells were harvested and pellets were lysed and prepared via boiling for 15 mins in 1X loading dye (50 mM Tris [pH 6.8], 100 mM DTT, 2% SDS, 0.1% bromophenol blue, 10% glycerol). Lysates were separated by SDS-PAGE on 10% Bis-Tris Plus Bolt gels (ThermoFisher) and transferred onto 0.45μm nitrocellulose membrane. Membranes were blocked with 2% nonfat dry milk in 1X TBST (10 mM Tris [pH 8], 150 mM NaCl, 0.05% Tween-20), and detection of indicated proteins was carried out using the following antibodies: Aiolos (clone D1C1E, Cell Signaling, 1:20,000), Bcl-6 (clone K112, BD Biosciences, 1:500), pSTAT5(Y694/9) (clone BD Biosciences, 1:5000), STAT5 (clone (D206Y, Cell Signaling, 1:5000), β-Actin (GenScript, 1:15,000), T-bet (clone 4B10 Santa Cruz, 1:500), Eomes (Abcam. 1:5,000) goat anti-mouse:HRP (Jackson Immunoresearch, 1:5,000-1:30,000), mouse anti-rabbit:HRP (Santa Cruz, 1:5000-1:20,000).

### RNA-seq Analysis

Naïve CD4^+^ T cells were cultured under T_H_1 or TFH-polarizing conditions for 3-4 days. Total RNA was isolated using the Macherey-Nagel Nucleospin kit according to the manufacturer’s instructions. Samples were provided to Azenta Life Sciences for polyA selection, library preparation, sequencing, and DESeq2 analysis. Genes with a p-value < 0.05 were considered significant, and those with an absolute log2 fold change of >1.0 were defined as differentially expressed genes (DEGs) for each comparison in the present study. Genes pre-ranked by multiplying the sign of the fold-change by −log_10_ (p-value) were analyzed using the Broad Institute Gene Set Enrichment Analysis (GSEA) software for comparison against ‘hallmark’, ‘gene ontology’, and ‘immunological signature’ gene sets. Heatmap generation and clustering (by Euclidean distance) were performed using normalized log2 counts from DEseq2 analysis and the Morpheus software (https://software.broadinstitute.org/morpheus). Volcano plots were generated using −log_10_(p-value) and log2 fold change values from DEseq2 analysis and VolcaNoseR software (https://huygens.science.uva.nl/VolcaNoseR/)^52^.

### Influenza infection and tissue preparation

Influenza virus strain PR8 was propagated in 10-day old embryonated chicken eggs and titered on MDCK cells. Naive mice were infected intranasally with 40 PFU PR8 suspended in 50uL sterile PBS. After 8 days, draining lymph nodes were harvested. For serum antibody analyses, whole blood was collected by cardiac puncture for serum isolation. Single cell suspensions were generated in tissue preparation media (IMDM + 4% FBS) by passing tissue through a nylon mesh strainer, followed by erythrocyte lysis via 3-minute incubation in 0.84% NH_4_Cl. Cells were washed 1X in ice cold FACS buffer (PBS + 4% FBS) before staining.

### Human Tissue Samples

Human tissues were collected and utilized in accordance with protocols approved by The Ohio State University Institutional Review Board. Donor consent was acquired when deemed appropriate according to the approved Ohio State University Institutional Review Board protocol. Human pediatric tonsils were obtained following overnight shipment after surgery via the CHTN Western Division at Vanderbilt University (Nashville, TN). Lymphocytes were enriched from fresh tonsil tissue specimens using previously reported protocols^53^. Briefly, single-cell suspensions were generated by dissociation via a GentleMACS Dissociator (Miltenyi Biotech) according to the manufacturer’s instructions. Cells were diluted in PBS (Thermo Fisher Scientific), layered over Ficoll-Paque PLUS (GE Healthcare), and centrifuged at 2000 rpm for 20 minutes at room temperature with the brake off and the mononuclear layers were harvested.

### Flow Cytometry

For analysis of antigen-specific CD4^+^ T cell populations in murine samples, cells were stained in FACS buffer with IA^b^ NP_311–325_ MHC class II tetramer (1:100, NIH Tetramer Core Facility) at room temperature for 1 hour. To address intra-replicate variability for the analysis of antigenspecific populations, the percentage of NP^+^ cells in Aiolos-deficient samples was presented relative to WT. For these analyses, a single WT sample was set as the control for each independent experiment, and each sample was normalized to this control. For the analysis of extracellular markers, samples were pre-incubated for >5 minutes at 4C with Fc Block (clone 93; BioLegend), then stained in the presence of Fc block for 30 minutes at 4C using the following antibodies: CD4 (1:300; clone GK1.5; R&D Systems); CD44 (1:300; clone IM7; BD Biosciences); CD62L (1:300; clone MEL-14; ThermoFisher); PD-1 (1:50; clone 29F.1A12; BioLegend); Cxcr5 (1:50; clone SPRCL5; ThermoFisher); CD25 (1:25; clone PC61.5; ThermoFisher); Cx3cr1 (1:100; Clone Z8-50 (RUO); BD Biosciences), and Ghost viability dye (1:400; Tonbo Biosciences). Cells were then washed 2X with FACS buffer prior to intracellular staining. For intracellular staining, cells were fixed and permeabilized using the eBioscience Foxp3 transcription factor staining kit (ThermoFisher) for 30 minutes at room temperature, or overnight at 4C. Following fixation, samples were stained with the following antibodies in 1X eBiosciences permeabilization buffer for 30 minutes at room temperature: T-bet (1:100; clone 4B10; BioLegend); Foxp3 (1:300; clone FJK-16s; ThermoFisher); Bcl-6 (1:20; clone K112-9; BD Biosciences); Eomes (1:100; clone DAN11MAG; ThermoFisher); Granzyme B (1:100; clone GB11 ThermoFisher); and Perforin (1:100; S16009A; BioLegend). Cells were washed with 1X eBiosciences permeabilization buffer and resuspended in FACS buffer for analysis. Samples were run on a BD FACS Canto II and analyzed using FlowJo software (version 10.8.1). For human tissue analysis, lymphocyte populations were stained using antibodies directed against surface or intracellular proteins according to the manufacturers’ recommendations. Where appropriate, fluorescence minus one (FMO) controls were used to determine nonspecific staining. The LIVE/DEAD Fixable Aqua Dead Cell Stain Kit (Thermo Fisher Scientific) was used to exclude nonviable cells in the analysis. Intracellular staining was performed using the Transcription Factor/FOXP3 Fixation/Permeabilization Solution Kit (Thermo Fisher Scientific) according to the manufacturer’s instructions. For surface staining, the following antibodies were used: CD44-APC (DB105, Miltenyi Biotec), CD4-AF700 (RPA-T4, Thermo Fisher Scientific), CD45RA-APC-cy7 (REA1047, Miletnyi Biotec), AIOLOS-PE (16D9C97, BioLegend), CXCR5-Pe-Vio770 (MU5UBEE, Thermo Fisher Scientific), FOXP3-PerCP-Vio770 (PCH101, Thermo Fisher Scientific), CD3-PE-Vio615 (UCHT1, BioLegend), CD8-FITC (REA734, Miltenyi Biotec), PD-1-BV421 (MIH4, BD Biosciences), CD62L-BV605 (DREG-56, BD Biosciences), CD45RO-BV650 (UCHTL1, Biolegend). Data were acquired on the FACSAria II (BD Biosciences) and analyzed using FlowJo (BD Biosciences) software.

### Serum antibody ELISA

To detect anti-PR8 antibodies response, high-binding microplates (Corning) were coated with 50 μL of UV-inactivated PR8 in PBS at 4°C overnight. To quantify the absolute antibody concentration, a standard curve for mouse isotype was established for each plate by coating 2 columns with 50 μL goat anti-mouse Ig (1 μg/mL in PBS; Southern Biotech) overnight at 4°C. Plates were blocked with 200 μL 1% bovine serum albumin (Fisher Scientifics) in PBSTE (Phosphate buffered saline with 0.05% Tween-20 and 1mM EDTA) for 1 hour at 37°C. After washing with PBSTE, 50 μL of samples and standard Ig (Southern Biotech) were added to respective wells and incubate for 1 hour at room temperature (RT). For total IgG standard, a mixture of purified mouse IgG1, IgG2b, IgG2c, and IgG3 were combined to emulate WT antibody titers^54^. Biotinylated goat-anti-mouse isotype-specific (IgM, IgG) antibodies (50 μL at 0.1 μg/mL; Southern Biotech) were added and incubated at RT for 1 hour. Horseradish-peroxidase-conjugated streptavidin (50 μL of 1:5000 dilution; ThermoFisher) was added and incubated at RT for 30 minutes. The plates were washed with PBSTE between steps for the procedures above. TMB High Sensitivity Substrate (BioLegend) was used for detection; reactions were stopped by adding 1 M sulfuric acid. Absorbances were measured at OD450 with OD540 as background using M2e SpectraMax 500 (Molecular Device) and antibody concentrations were calculated using the generated standard curve.

### ATAC-seq Analysis

ATAC-seq analysis was performed as described^55^. Briefly, 5×10^4^ cells at >95% viability were processed using the Illumina Nextera DNA Library Preparation Kit, according to the manufacturer’s instructions. Resultant sequences were trimmed using Bowtie 2 and trimmed reads were aligned to mouse genome mm10. Statistically significant peaks were identified using MACs2. CPM-normalized tracks were visualized using Integrative Genomics Viewer (IGV) version 2.12.2. For all presented loci, CPM ranges (0.5-2.0) are equivalent between genotypes.

### Chromatin Immunoprecipitation (ChIP)

ChIP assays were performed as described previously^51^. Resulting chromatin fragments were immunoprecipitated with antibodies against Aiolos (Cell Signaling, clone D1C1E, 2ug/IP), STAT5 (R&D AF2168, 5ug/IP) or IgG control (Abcam ab6709; 2-5ug/IP, matched to experimental antibody). Enrichment of the indicated proteins was analyzed via qPCR with the primers below. Samples were normalized to total DNA controls and isotype control antibodies were used to ensure lack of non-specific background.

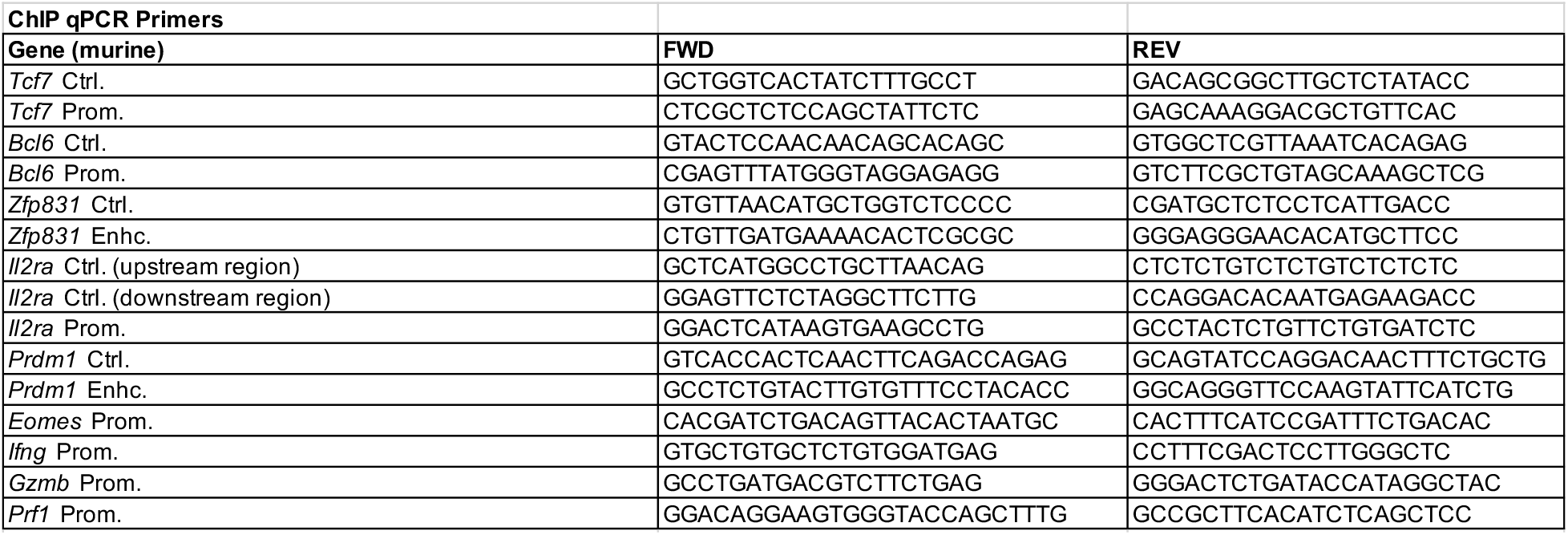

### Statistical analysis

All statistical analyses were performed using the GraphPad Prism software (version 9.3.1). For single comparisons, unpaired Student’s Wests were performed. For multiple comparisons, oneway ANOVA with Tukey’s multiple comparison tests were performed. Error bars indicate the standard error of the mean. *P* values <0.05 were considered statistically significant.

## Supporting information

Supplementary Data Figures 1 through 6

## Acknowledgments

The authors would like to thank all members of the Oestreich Lab, as well as colleagues in the Department of Microbial Infection and Immunity for constructive criticism. The authors would also like to thank the Cooperative Human Tissue Network of the Nationwide Children’s Hospital (Columbus, OH) for providing human pediatric tonsil samples. The authors would like to thank members of the Lio laboratory (Heng-Yi Chen and Fang-Yun Lay) for their technical advice. Finally, the authors would like to thank members of the Yount laboratory (Ashley Zani, Adam Kenney, and Lizhi Zhang) for assistance with influenza virus preparation.

## Author contributions

D.M.J. and K.A.R. assisted with the design of the study, performed experiments, analyzed data, and wrote the manuscript. S.P., J.A.T., E.D.S.H., A.V., C.D.E., R.T.W., and A.G.F. performed experiments and analyzed data. P.L.C., M.D.P., and J.M.B. assisted with analysis of -omics data. A.S., O.A., E.A.H, J.S.Y., H.E.G., and C.J.L. assisted with the influenza infection experiments and analyzed data. K.J.O. supervised the research, designed the study, analyzed data, and wrote the manuscript.

