## Supplementary Data Figures 1 through 6 for "Aiolos modulates the T_FH_ and CD4-CTL differentiation programs via reciprocal regulation of the Zfp831/TCF-1/Bcl-6 axis and CD25"

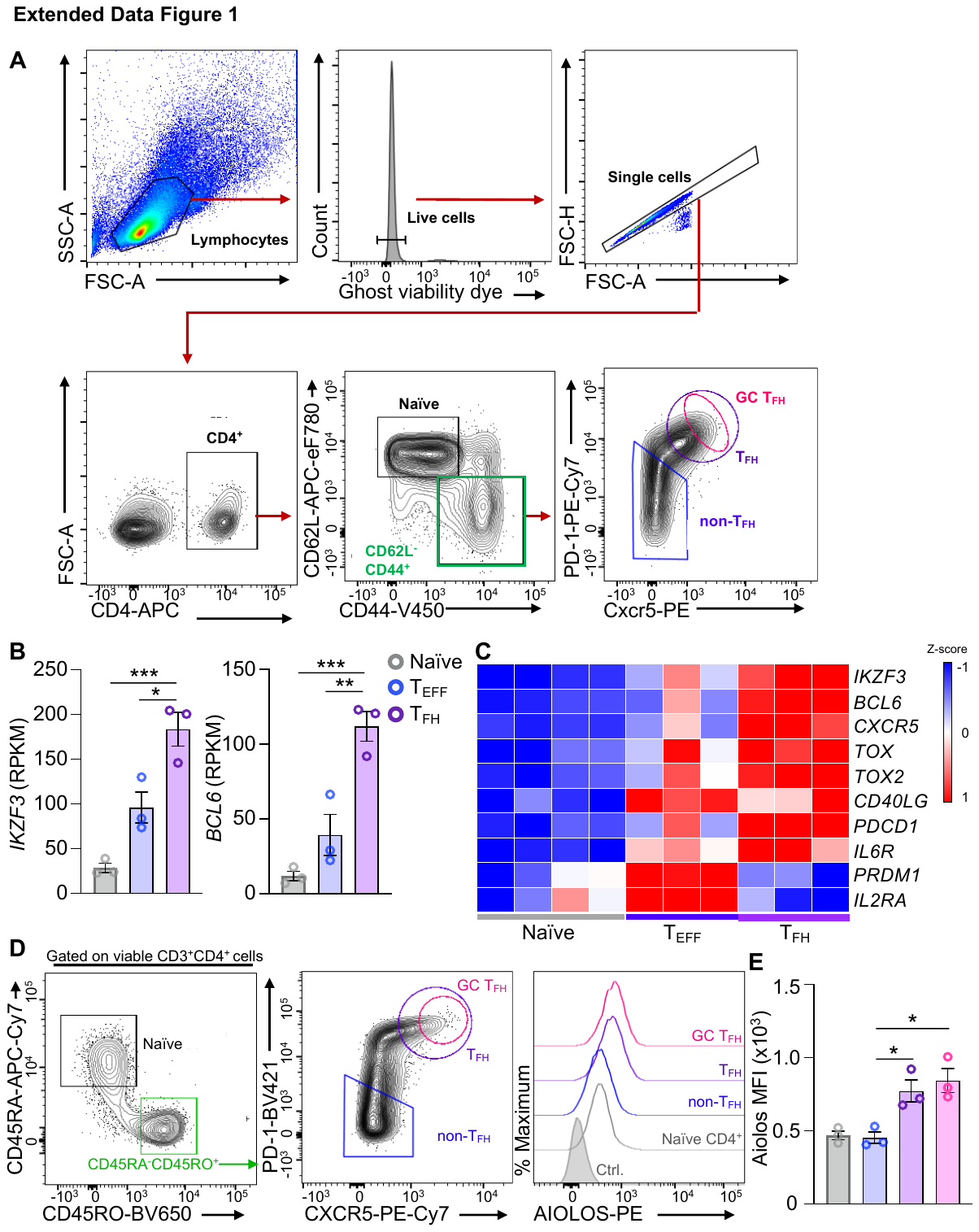

**Extended Data Figure 1.** (A) Gating strategy utilized for analysis of CD4^+^ T cell populations generated during influenza infection. Example gating for T_FH_ populations is shown. **(B-C)** Publicly available RNA-seq data (GEO# GSE58596) from human tonsillar CD4^+^ naïve, effector (T_EFF_), and T_FH_ populations. Data are presented as reads per kilobase of transcript per million mapped reads (RPKM) (B) or as a heatmap with reads normalized by row (gene) (C). Data are representative of 3 biological replicates (n = 3 ± s.e.m; **P* < 0.05, ***P* < 0.01, ****P* <0.001; one-way ANOVA with Tukey’s multiple comparison test). **(D,E)** Representative flow cytometry data of AIOLOS expression in human CD4^+^ T cell populations isolated from pediatric tonsils. Median fluorescence intensity was quantitated for naïve (CD3^+^CD4^+^CD8^-^CD45RA^+^CD45RO^-^; grey bar), non-T_FH_ effector (CD3^+^CD4^+^CD8^-^CD45RA^-^CD45RO+PD-1^lo^CXCR5^-^; blue bar), T_FH_ (CD3^+^CD4^+^CD8^-^CD45RA^-^CD45RO^+^PD-1^hi^CXCR5^+^; purple bar), and germinal center (GC) T_FH_ (CD3^+^CD4^+^CD8^-^CD45RA^-^CD45RO^+^PD-1^hi^/CXCR5^hi^; purple bar) populations. Data are representative from two independent experiments (n = 3 ± s.e.m; **P* < 0.05; unpaired Student’s t-test).

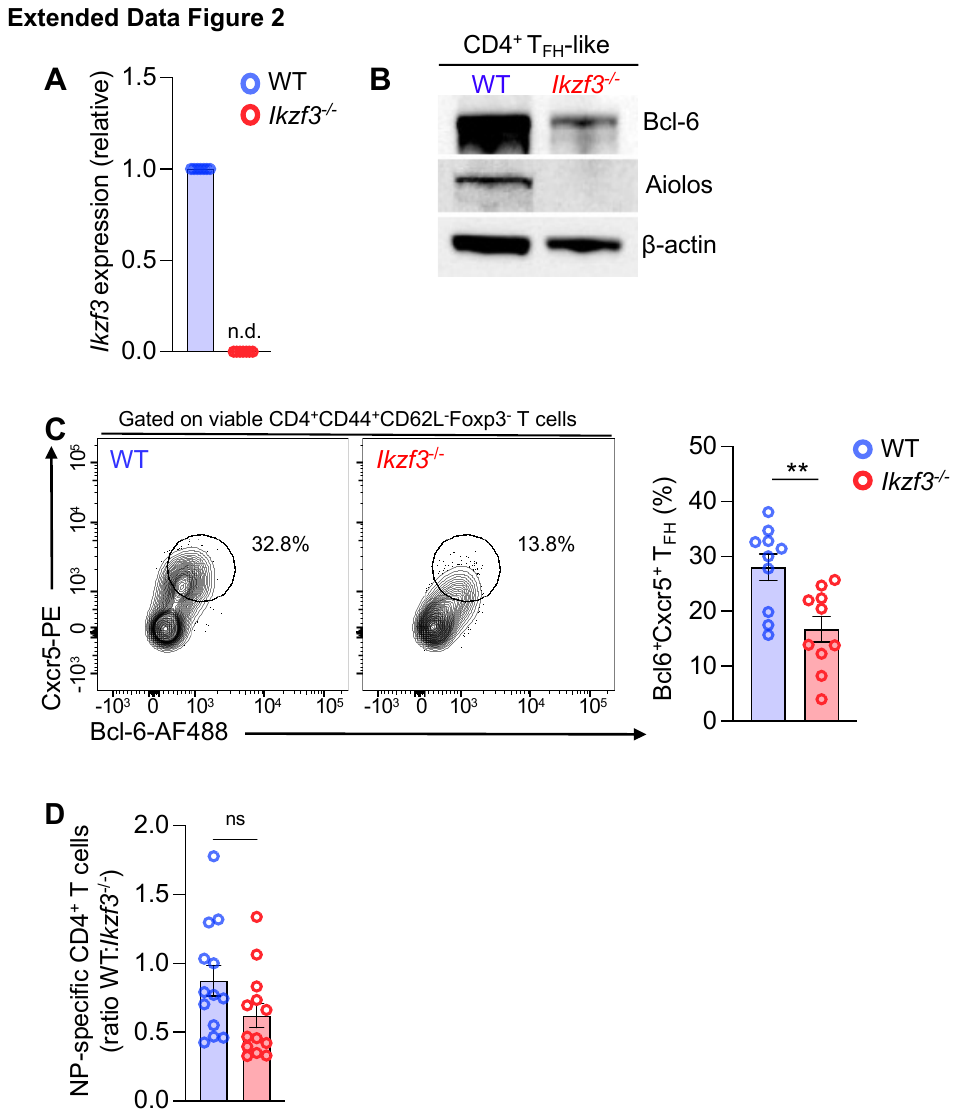

**Extended Data Figure 2. (A-B)** Naïve WT or Aiolos-deficient (*Ikzf3*^-/-^) CD4^+^ T cells were cultured under T_FH_-polarizing conditions for 3 days. (A) Analysis of Aiolos transcript was performed via qRT-PCR. (n=7 ± s.e.m; n.d., not detected). (B) Analysis of the indicated proteins was performed via immunoblot. B-actin serves as a loading control. Image is representative from 2 biological replicates from 2 independent experiments. **(C-D)** Naïve WT or *Ikzf3*^-/-^ C57BL/6 mice were infected intranasally with 40 PFU influenza (A/PR8/34; “PR8”) for 8 days. (A). (C) The percentage of bulk Bcl-6^HI^Cxcr5^HI^ T_FH_ populations was assessed. Data are compiled from 3 independent experiments (n=10 ± s.e.m; ***P* < 0.01; unpaired Student’s t-test). (D) Ratio of total numbers of NP-specific CD4^+^ T cells. To account for intra-replicate variability, the percentages of NP^+^ cells (WT and *Ikzf3*^-/-^) are presented relative to a single WT control per experiment. Data are compiled from 4 independent experiments. (n=13 ± s.e.m; unpaired Student’s t-test).

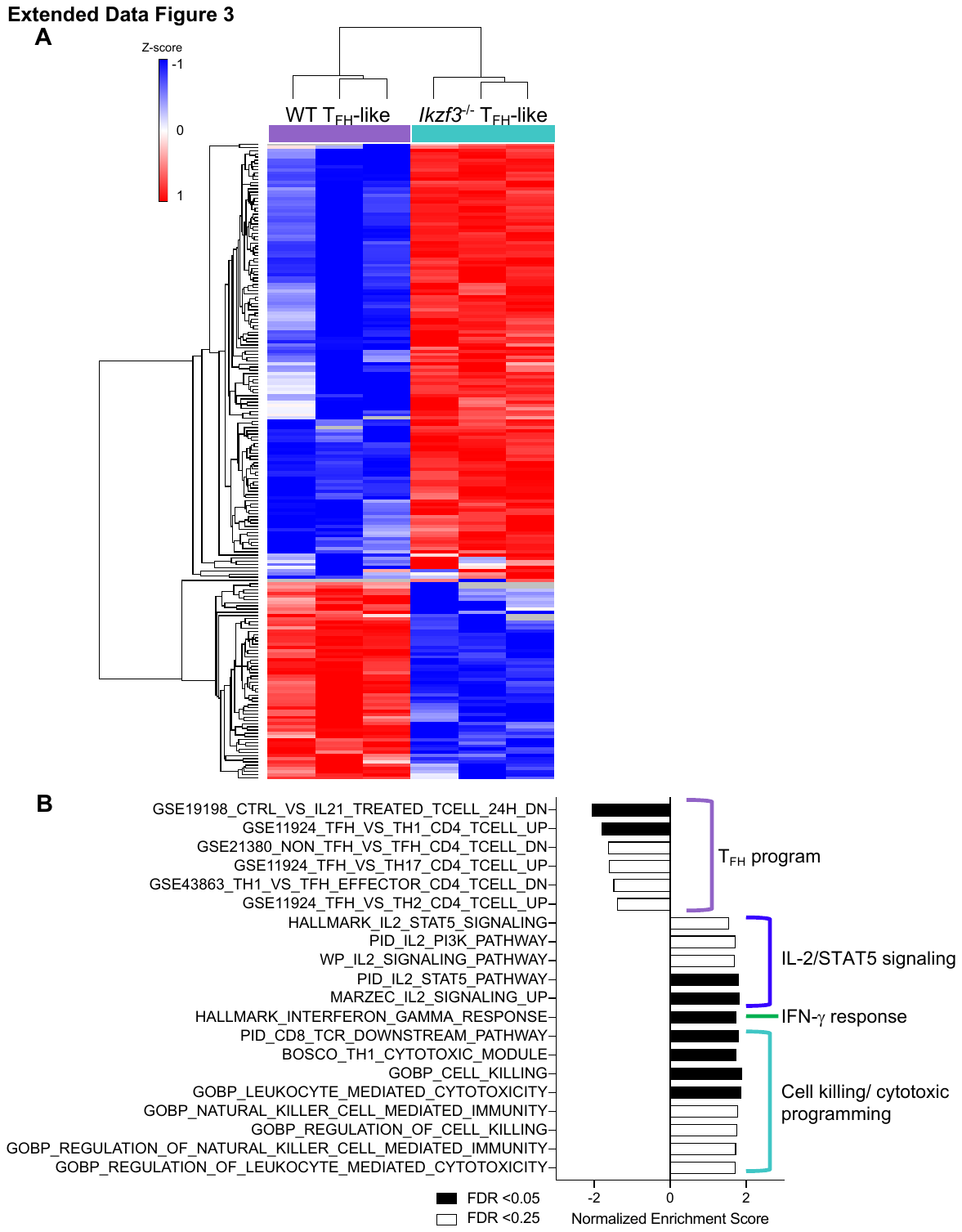

**Extended Data Figure 3.** Naïve WT or Aiolos-deficient (*Ikzf3*^-/-^) CD4^+^ T cells were cultured under T_FH_-polarizing conditions for 3 days. RNA-seq analysis was performed to assess differentially expressed genes (DEGs) between WT and Aiolos-deficient cells. (**A**) Heatmap displaying the top 200 DEGs. Clustering was performed using Euclidean distance. Data are compiled from 3 biological replicates from 3 independent experiments. (**B**) GSEA analysis of pre-ranked DEGs, compared against ‘hallmark’, ‘curated’, ‘immunological signature’, and ‘gene ontology’ gene sets. Data are compiled from 3 biological replicates from 3 independent experiments.

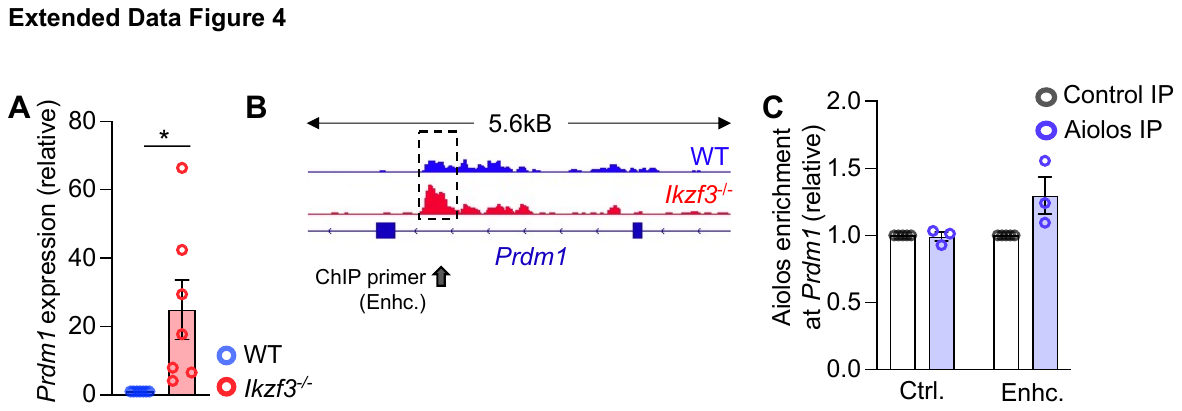

**Extended Data Figure 4.** Naïve CD4^+^ T cells were cultured under T_FH_-like polarizing conditions for 3 days. (**A**) qRT-PCR analysis of *Prdm1* transcript. Data were normalized to *Rps18* control and are presented relative to the WT sample. (n=7 ± s.e.m; **P* < 0.05; unpaired Student’s t-test). **(B)** Representative ATAC-seq analyses of the *Prdm1* locus from two independent experiments are displayed as CPM-normalized Integrative Genomics Viewer (IGV) tracks. Sites of altered accessibility, and approximate ChIP primer locations, are indicated. **(C)** ChIP analysis of Aiolos enrichment at the indicated region, or negative control region, are shown. Data were normalized to total IgG and presented relative to enrichment in the IgG IP control sample. (n=3 ± s.e.m).

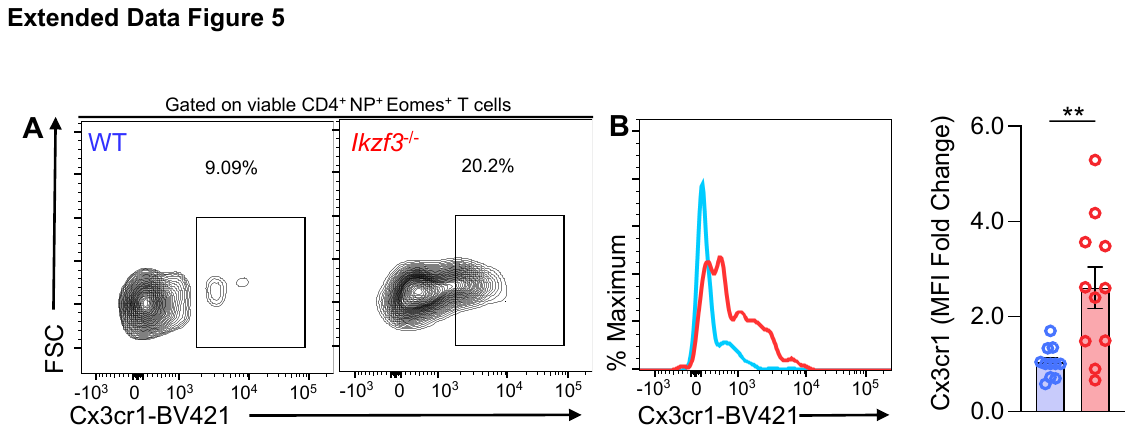

**Extended Data Figure 5.** Naïve WT or *Ikzf3*^-/-^ C57BL/6 mice were infected intranasally with 40 PFU influenza (A/PR8/34; “PR8”) for 8 days. (A) Analyses of the percentage of antigen-specific (NP^+^) Cxcr3r1^+^ cells were performed via flow cytometry. (B) Histogram analysis of Cx3cr1 and fold change in median fluorescence intensity. To account for intra-replicate variability, the percentage of NP^+^Cx3cr1^+^ cells in Aiolos-deficient samples was presented relative to WT. For these analyses, a single WT sample was set as the control for each independent experiment, and each sample was normalized to this control. Data are compiled from 4 independent experiments (n=11 ± s.e.m; ***P* < 0.01; unpaired Student’s t-test).

**Extended Data Figure 6.** Naïve CD4^+^ T cells were cultured under T_H_1 polarizing conditions for 3 days. (**A**) qRT-PCR analysis of *Il2ra* and *Prdm1* transcript. Data were normalized to *Rps18* control and presented relative to the WT sample. (n=9 ± s.e.m; ***P* < 0.01, ***<0.001; unpaired Student’s t-test). **(B)** Representative ATAC-seq analyses of the indicated loci from two independent experiments are displayed as CPM-normalized Integrative Genomics Viewer (IGV) tracks. Sites of altered accessibility, and approximate ChIP primer locations, are indicated.
